# The antibacterial factor APOL3 couples lysosomal damage to mitochondrial DNA efflux and type I IFN induction

**DOI:** 10.1101/2025.05.16.654477

**Authors:** Dominic A. Ritacco, Hamna Shahnawaz, Antonia Oduguwa, Jacob Hawk, Brianna Vizcaino, Donna Farber, Ryan G. Gaudet

**Affiliations:** Department of Microbiology and Immunology, Columbia University Irving Medical Center, New York, NY; Department of Surgery, Columbia University Irving Medical Center, New York, NY

## Abstract

Lysosomal damage is an endogenous danger signal to the cell, but its significance for innate immunity and how specific signaling pathways are engaged by this stressor remain unclear. Here, we uncover an immune-inducible pathway that connects lysosomal damage to mitochondrial DNA (mtDNA) efflux and type I IFN production. Lysosomal damage elicits mitochondrial outer membrane permeabilization (MOMP) via BAK/BAX macropores; however, the inner mitochondrial membrane (IMM) prevents wholesale mtDNA release in resting cells. Priming with type II IFN (IFN-γ) induced the antibacterial effector apolipoprotein L-3 (APOL3), which upon transient lysosomal damage, targets mitochondria undergoing MOMP and selectively permeabilizes the IMM to enhance mtDNA release and activate cGAS/STING signaling. Biochemical and cellular reconstitution revealed that analogous to its bactericidal detergent-like mechanism, APOL3 solubilizes cardiolipin to permeabilize the IMM. Our findings illustrate how cells use an antibacterial protein to expedite the breakdown of endosymbiosis and facilitate a heightened response to injury and infection.

## INTRODUCTION

Lysosomes are vulnerable to damage by the material they transport. As the endpoint of endocytosis, phagocytosis and autophagy, these organelles are continually threatened with a barrage of infectious (bacteria, viruses) and non-infectious (particulates, crystals, toxins) insults. Rupture of the enclosing membrane releases a harmful cocktail of acidic hydrolases into the cell and is a hallmark of infection, inflammatory sequelae, neurodegeneration, and cancer^1^. Widespread lysosomal destabilization triggers a variety of cell death pathways including as apoptosis, necroptosis and pyroptosis^2–4^. Cells have thus evolved many ways of repairing or removing damaged lysosomes^5–10^. Transient lysosomal damage can also have pronounced effects on cellular homeostasis, and is perceived by the innate immune system as an endogenous danger signal^11^. Signaling outputs emanating from sub-lethal lysosomal perturbation include inflammasome activation^12^, interferon (IFN) signaling^13–15^, chromosome segregation^16^, antigen presentation^17,18^ and metabolic reprogramming^19^. However, the mechanisms by which lysosomal damage triggers such disparate signaling pathways in specific contexts is poorly understood.

Damaged lysosomes communicate with other organelles like mitochondria according to the scale of injury ^19–22^. Severe lysosomal damage activate proapoptotic BAK and BAX to initiate mitochondrial outer membrane permeabilization (MOMP) and apoptosis ^3,23–27^; whereas transient damage reprograms mitochondrial metabolism^19^. Beyond these essential roles in energy metabolism and cell death, mitochondria have also emerged as innate immune signaling hubs. This occurs primarily through the regulated release of damage associated molecular patterns (DAMPs) including cytochrome *c*, reactive oxygen intermediates, and nucleic acids – each of which initiates signaling by a distinct pathway with disparate cellular effects ^28,29^. Though MOMP has historically been considered the point of no return for apoptosis, there is compelling evidence that MOMP can also occur in living cells^30^, and mitochondrial DAMPs are known to regulate cellular outputs to sub-lethal stressors like infection^31^ and senesence^32^. Thus, an intriguing hypothesis is that transient lysosomal damage is transmitted to mitochondria to integrate and transduce a selective cellular response.

In contrast to type I IFN (IFN-α/β) which can be made by most cell types and induces antiviral interferon-stimulated genes (ISGs), type II IFN (IFN-γ) is produced by lymphocytes and elicits expression of broadly antimicrobial genes^33^. One group of effectors that are strongly induced by IFN-γ are a cluster of six poorly understood primate-specific genes termed the Apolipoprotein L (APOL) family ^34,35^. We previously discovered *APOL3* encodes an intracellular innate immune restriction factor that coats cytosol-invasive bacteria like *Salmonella enterica* serovar Typhimurium and mediated bacteriolysis by dissolving the inner bacterial membrane into discoidal lipoprotein particles^36^. Interestingly, APOL3 was not a bacterial specific receptor, instead it was initially recruited to host endomembrane structures damaged by bacteria. Indeed, sterile insults were sufficient to recruit APOL3 to leaky lysosomes^36^. Despite this, a role for APOL3 in the response to lysosomal damage beyond bacterial restriction has not been identified. Moreover, whether immune cytokines like IFN-γ recalibrate the way target cells react to organellar injury in general is poorly understood.

Here, we demonstrate that in living cells, APOL3 couples lysosomal damage to mtDNA efflux and type I IFN production. Transient lysosomal damage inflicted by sterile or infectious stressors triggered APOL3 recruitment to BAX/BAK-permeabilized mitochondria where it selectively enhanced IMM permeability to mtDNA by solubilizing cardiolipin. We present evidence that transient lysosomal damage induces sub-lethal MOMP; however, in resting cells, IMM integrity maintains a barrier against wholesale mtDNA release. IFN-γ priming elicited APOL3 which negated this barrier by permeabilizing the IMM, thereby facilitating a physical interaction between mtDNA and cGAS to induce type I IFNs without initiating cell death. Our results illustrate how cells use an immune-inducible antibacterial protein to expedite the selective breakdown of endosymbiosis and facilitate a heightened response to injury and infection.

## RESULTS

### APOL3 mediates cGAS-dependent type I IFN signaling downstream of lysosomal damage

We set out to investigate if the antibacterial effector APOL3 played a role in the cellular response to sterile lysosomal damage. HeLa cells harboring a CRISPR/Cas9 deletion in the *APOL3* locus or isogenic non-targeting control (NTC) cells cultured in parallel were primed overnight with IFN-γ to induce *APOL3* expression, then treated with a sub-lethal dose of the lysosomotropic agent Leucyl-L-leucine methyl ester (LLOME) which is commonly used to rupture endolysosomes^5,10,37^. Analysis of Galectin-3 as marker for damaged lysosomes, or Magic Red® fluorescence as a readout of lysosomal cathepsin activity, revealed *APOL3* deletion did not affect the number of lysosomes permeabilized by LLOME nor did it effect the speed of lysosomal recovery following removal of LLOME (Figure S1A and S1B). Cells possess multiple mechanisms to repair lysosomal damage ^5–9^, but whether transient lysosomal injuries regulate other cellular pathways downstream of the initial damage is less understood. To address this, we took an unbiased approach to analyze the transcriptional response in NTC or APOL3^-/-^ cells after recovery from lysosomal damage. After priming overnight with IFN-γ, cells were pulsed with LLOME, then allowed to recover for 8 h before RNAseq analysis. Surprisingly, differential expression between LLOME treated and untreated NTC cells revealed the most prominent genes upregulated by lysosomal damage were a subset of antiviral ISGs including *MX1, IFIT3*, *RSAD2* and *OASL* (Figure 1A). A second set of stress-related genes (*IER3*, *ATF3*, *DUSP5*, *JUN*, *ANKRD1*, *MAFF*) was also upregulated by LLOME (Figure 1A). This latter set was more expected given that many have been identified by others as upregulated by LLOME in resting cells^38^. Strikingly, LLOME-triggered upregulation of ISGs was severely blunted in APOL3^-/-^ cells whereas induction of stress response was intact and comparable to that observed for NTC cells (Figure 1A). Indeed, comparing LLOME-treated NTC and APOL3^-/-^ cells identified downregulation of a type I IFN signature as the predominant differentially expressed gene set (Figure S2A). In contrast, no differentially expressed genes were identified between IFN-γ-primed NTC and APOL3^-/-^ cells in mock treated samples (Figure S2B). qPCR for individual transcripts confirmed that both APOL3 and IFN-γ-priming were essential for LLOME-triggered upregulation of antiviral ISGs but were dispensable for the induction of stress response genes (Figure S2C, and S2D). These data suggest that for IFN-γ-primed cells, transient lysosomal damage triggers selective upregulation of antiviral ISGs in a manner that requires APOL3.

**Figure 1.**
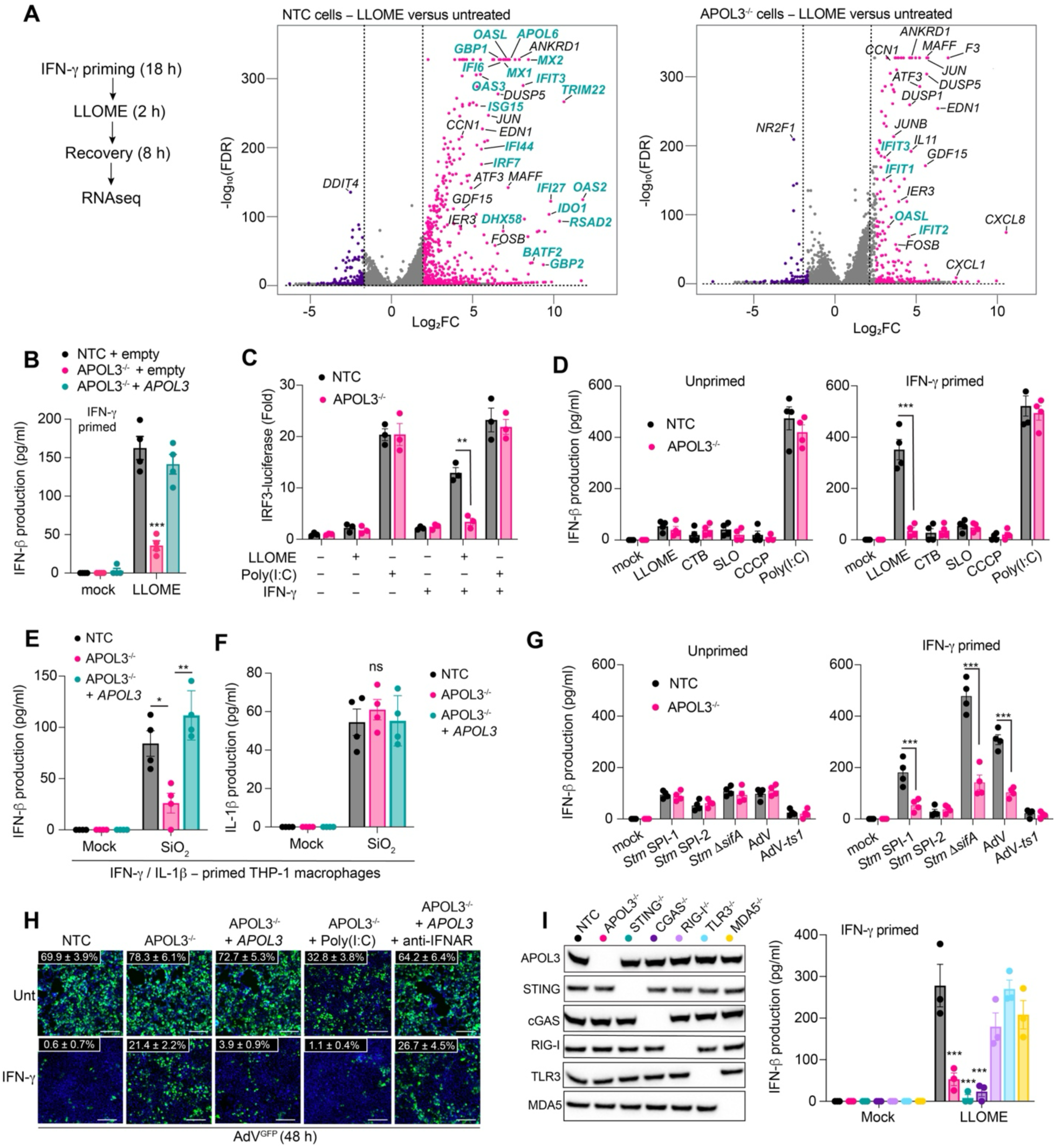
Transient lysosomal damage elicits APOL3 and cGAS-dependent type I IFN signaling. (A) NTC or APOL3^-/-^ HeLa cells primed overnight with IFN-γ, were pulsed with LLOME (600 μM, 2 hr unless otherwise indicated), recovered for 8 h and analyzed by RNAseq. Volcano plots show differential expression between untreated (DMSO) and LLOME treated for both genotypes. Significant genes (P < 0.01, log_2_-fold change (LFC) > 2) are depicted in rose (upregulated) or purple (downregulated). Selected genes of interest were annotated (ISGs in teal). Statistical significance was calculated using the Wald test, with P values adjusted by the Benjamini-Hochberg procedure. (B) NTC or APOL3^-/-^ HeLa cells expressing pMSCV-empty or pMSCV-*APOL3* were pulsed with LLOME and IFN-β protein levels in the supernatants determined by ELISA after 12 hr. (C) HeLa genotypes expressing IRF3-luciferase +/- IFN-γ priming pulsed with LLOME or treated with Poly(I:C) and luminescence measured after 6 hr. Data is fold change relative to mock treated. (D) IFN-β in the supernatants of primed or unprimed NTC or APOL3^-/-^ HeLa cells after the indicated insult. (E and F) IFN-β (E) or IL-1β (F) in the supernatants of IFN-γ/IL-1β-primed THP-1 macrophages 24 hr after being fed SiO_2_ nanoparticles (50 μg/ml for 4 hr) to trigger phagosomal rupture. APOL3 was expressed in *trans* via lentivirus transduction. (G) IFN-β levels in the supernatants of NTC or APOL3^-/-^ cells infected (8 hr) with invasive (SPI-1), hyper-invasive (Δ*sifA*), or non-invasive (SPI-2) *Salmonella* Typhimurium (*Stm*), invasive adenovirus (AdV), or non-invasive AdV-*ts1*. (H) HeLa genotypes +/- IFN-γ priming infected with AdV-GFP (24 hr) and imaged (representative images shown). Poly(I:C) or anti-IFNAR antibodies were co-incubated for the duration. Percentage values denote mean ± SD, *n* = 3) (I) IFN-β in the supernatants of IFN-γ-primed HeLa cells of the indicated genotypes 24 hr after LLOME pulse. Representative immunoblots for each genotype are shown. Data represent mean ± SEM from 3 or 4 independent experiments. *P < 0.05; ** P < 0.01; *** P < 0.001 determined by one-way ANOVA with Tukey’s multiple comparison test.

Notably, the suite of antiviral ISGs upregulated by LLOME are classic targets of type I IFN that were not induced by type II IFN (IFN-γ) priming alone (Figure S2D). Thus, we asked if lysosomal damage triggered APOL3-dependent type I IFNs that could induce these antiviral ISGs through paracrine or autocrine signaling. Transient LLOME exposure resulted in significant IFN-β protein secretion by IFN-γ-primed NTC cells but not APOL3^-/-^ cells; an effect that was rescued by expression of APOL3 in *trans* (Figure 1B). Looking upstream at the IRF3 transcription factor that promotes *IFNB1* transcription^39^, LLOME induced APOL3-dependent IRF3-luciferase activation but only in IFN-γ-primed cells wherein APOL3 is expressed (Fig. 1C). This is consistent with our previous findings that *APOL3* is specifically induced by IFN-γ and not IFN-α or IFN-β in HeLa cells^36^. Notably, IRF3 induction by the TLR3 agonist Poly(I:C) was comparable between NTC and APOL3^-/-^ cells implying the IRF3 signaling pathway remained intact after APOL3 deletion (Figure 1C). To further investigate the specificity of this response, we damaged other organelles and measured IFN-β production with or without priming. Relative to LLOME, damaging the Golgi apparatus (cholera toxin-β), the plasma membrane (streptolysin-O), or mitochondria (carbonyl cyanide m-chlorophenyl hydrazone (CCCP)) induced much lower IFN-β levels that were only slightly above background (Figure 1D). In a different cellular system, we examined cytokine production in THP-1 differentiated macrophages that were fed silicon dioxide (SiO_2_) nanoparticles to induce sub-lytic phagosomal disruption^5^. Here, overnight priming with two signals, IFN-γ and IL-1β, was required to elicit APOL3 protein expression (Figure S3A) and mediated maximal IFN-β secretion after SiO_2_ feeding (Figure S3B). Deletion of APOL3 in THP-1 cells suppressed SiO_2_-triggered IFN-β release, whereas ectopic expression of APOL3 restored this effect (Figure 1E). Silica particles also activate the NLRP3 inflammasome through phagosomal disruption^12,40^. However, APOL3^-/-^ cells produced normal levels of mature IL-1β after SiO_2_ treatment (Figure 1F). Thus, diverse sources of sterile lysosomal damage activate an APOL3-dependent type I IFN response in immune and non-immune cells.

We next asked whether this same process could be operative during infectious lysosomal damage. *Salmonella enterica* serovar Typhimurium (*Stm*) induces a type I IFN response in a CGAS/STING-dependent manner^41^. Whether endolysosomal damage is the triggering event and if APOL3 plays a role in this response has not been examined. In IFN-γ-primed cells, *Stm* induced an APOL3-dependent increase in IFN-β production when bacteria were cultured in SPI-I endomembrane-penetrating conditions but not when cultured in SPI-2 conditions that confine *Stm* to its entry vacuole (Figure 1G). Moreover, an *Stm* mutant (*sifA*) that induces more severe lysosomal damage triggered increased APOL3-dependent IFN-β production (Figure 1G). This response was independent of pathogen structure because adenovirus (AdV5), which also breach the endosome upon cell entry^42^, triggered a potent IFN-γ and APOL3-dependent IFN-β response whereas a temperature sensitive mutant (AdV-*ts1*) incapable of membrane penetration failed to induce IFN-β (Figure 1G). These bacterial and viral mutants confirm that lysosomal damage is the triggering event for APOL3-dependent type I IFN activation during infection.

Given the APOL3-depenent ISG signature included many antiviral ISGs, we asked if APOL3 conferred antiviral immunity by eliciting IFN-β production in response to lysosomal damage. Deletion of APOL3 suppressed IFN-γ-dependent restriction of AdV^GFP^, which was rescued by expression of APOL3 in *trans* or by activating an alternative IFN-β production with Poly(I:C) (Figure 1H). In contrast, neutralizing endogenous IFN-β signaling with anti-type I IFN receptor antibody abrogated APOL3-dependent AdV restriction (Figure 1H). Thus, APOL3 senses lysosomal damage and confers anti-viral immunity by potentiating type I IFN production.

Lastly, we considered the source of the type I IFNs triggered by lysosomal damage. We created a panel of CRISPR/Cas9 HeLa lines lacking canonical pattern recognition receptors that initiate IFN signaling – including toll-like receptor 3 (TLR3), cyclic GMP-AMP synthase (cGAS), stimulator of interferon genes (STING), retinoic acid-inducible gene I (RIGI) and melanoma differentiation-associated gene 5 (MDA5). IFN-γ-primed cells lacking cGAS or STING but not the other PRRs failed to secrete IFN-β when pulsed with LLOME, phenocopying APOL3 deletion (Figure 1I). These data suggest IFN-γ upregulates APOL3 to facilitate cGAS activation downstream of transient lysosomal damage.

### APOL3 relocates to mitochondria and facilitates mtDNA release during lysosomal damage

cGAS can be activated by cytosolic DNA of mitochondrial^43^ or nuclear^44^ origin. To ascertain the source of DNA that elicits APOL3-dependent cGAS activation during lysosomal damage, we immunoprecipitated FLAG-tagged cGAS overexpressed in IFN-γ-primed NTC or APOL3^-/-^ cells pulsed with LLOME and performed qPCR for genomic or mitochondrial DNA. Strikingly, LLOME-triggered selective cGAS binding to mtDNA but not abundant nuclear sequences like LINE1 elements or ribosomal DNA and this was severely blunted by deletion of APOL3 (Figure 2A). We then depleted mtDNA using 2′,3′-dideoxycytidine (ddC) in APOL3^-/-^ cells transduced with MSCV-*APOL3* and confirmed successful mtDNA depletion by qPCR (Figure S3C). In cells depleted for mtDNA, APOL3-dependent IFN-β-production triggered by sterile (LLOME) or infectious lysosomal damage was effectively suppressed to baseline levels (Figure 2B). Similar results were apparent for IFN-γ/IL-1β activated THP-1 macrophages where mtDNA depletion inhibited IFN-β induction by SiO2 particles (Figure S3D). We next assayed for cytoplasmic mtDNA in cells experiencing lysosomal damage to assess if APOL3 enhances mtDNA efflux from mitochondria. Lysosomal damage triggered APOL3-dependent mtDNA efflux specifically in IFN-γ primed cells (Figure 2C). Neither APOL3 deletion nor IFN-γ-priming had significant effect on mtDNA efflux triggered by the Bcl-2 inhibitor ABT-737 in conjunction with pan-caspase inhibition with Q-VD-OPh, (hereafter ABT-737/Q) (Figure 2C) which mediates BAK/BAX-dependent mtDNA release independently of lysosomal damage^10^. Thus, APOL3 enhances mtDNA efflux and cGAS activation specifically downstream of lysosomal damage.

**Figure 2.**
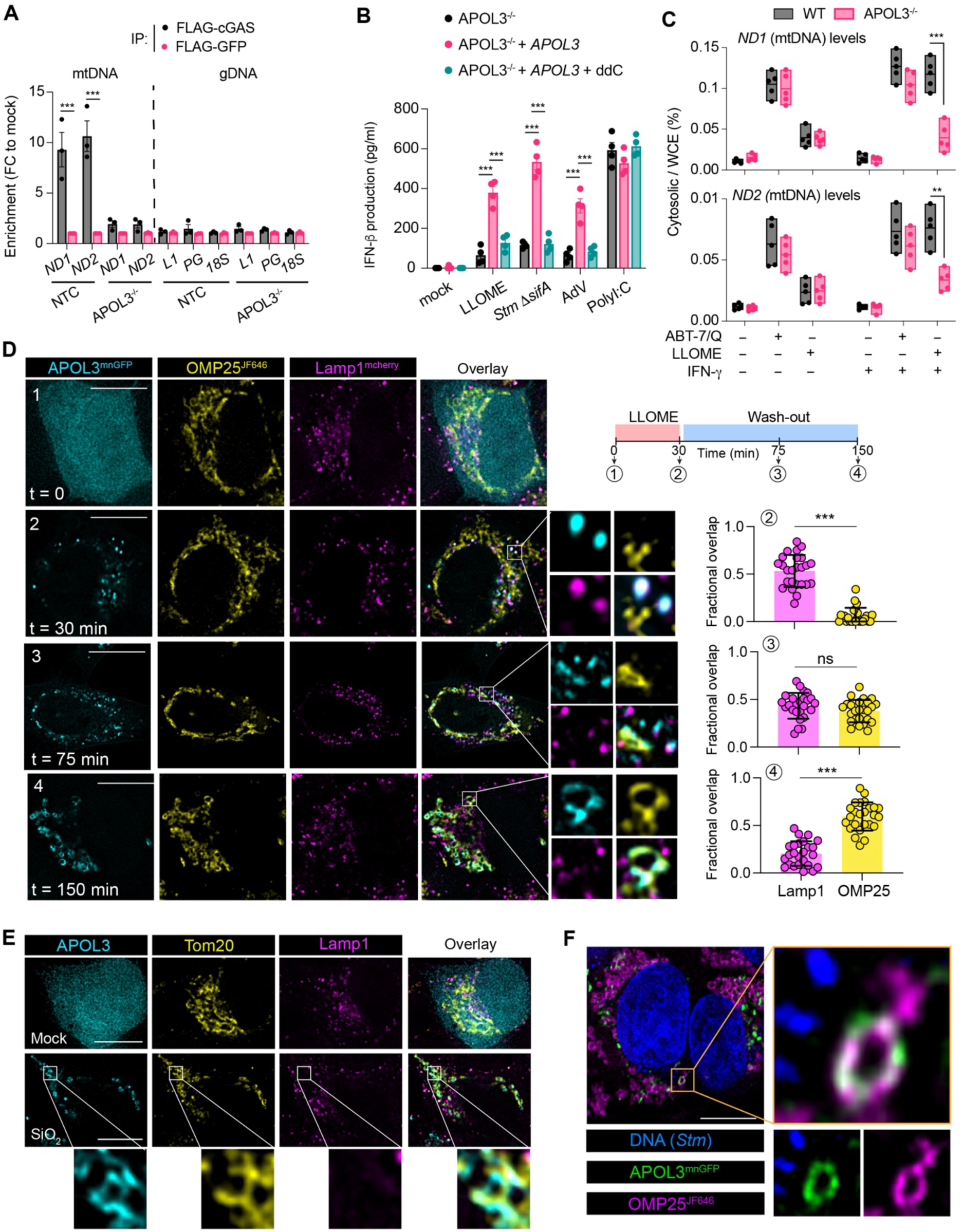
APOL3 forms puncta at mitochondria and enhances mtDNA release during lysosomal damage. (A) IFN-γ-primed HeLa genotypes expressing FLAG-cGAS or FLAG-GFP were treated with LLOME followed by extraction of DNA from FLAG immunoprecipitants and mitochondrial (*ND1*, *ND2*), nuclear (*L1*, *PG*) or ribosomal (*18S*) transcripts. Fold change (FC) is relative to mock treated. (B) IFN-β produced by IFN-γ-primed APOL3^-/-^ cells rescued with retroviral-*APOL3* and cultured with ddC to deplete mtDNA prior to treatment. (C) qPCR quantification of mtDNA in the cytosol relative to whole cell extract (WCE) after treatment with LLOME or ABT-737/Q. Shown are floating bars (mean, min/max, *n* = 5 independent experiments) (D) APOL3^mnGFP^ knock-in cells expressing OMP25^JF646^ and Lamp1-mCherry were pulsed with LLOME and imaged at the indicated time. Fractional overlap for APOL3 is shown (n = 25 cells, error bars ± SD). Results are representative of 3 independent experiments. (E) IFN-γ/IL-1β-primed THP-1 macrophages expressing APOL3-GFP were fed SiO_2_ particles (50 μg/ml, 3 hr) and immunostained for Tom20 or Lamp1. (F) APOL3^mnGFP^ cells infected with SPI-1 *Stm* (DAPI) for 3 hr and mitochondria labeled with OMP25^JF646^. Images are representative single *z* planes of Thunder deconvolved widefield images. Scale bar, 5 μm. Data are mean ± SEM from 3 (A) or 4 (B) independent experiments. *P < 0.05; **P < 0.01; ***P < 0.001 determined by two-tailed unpaired Student’s t tests (A, D) or one-way ANOVA with Tukey’s multiple comparison test (B, C).

These results prompted us to analyze subcellular location of APOL3. To study endogenous APOL3 dynamics, we edited the genomes of human epithelial cells to insert a C-terminal tag (mNeonGFP or HA) to the endogenous APOL3 protein (hereafter APOL3^mnGFP^ and APOL3^HA^ (Figure S4A). *APOL3* upregulation by IFN-γ was similar between wildtype (WT) and APOL3 knock-in HeLa cells (Figure S4B). Reaffirming results from overexpression studies^36^, APOL3^mnGFP^ was diffusely cytosolic at resting state but formed puncta at Lamp1+ lysosomes 30 min after exposure to LLOME (Figure S4C) and coated cytosol invasive (SPI-1) *Stm* 1 hr after infection (Figure S4D). However, APOL3 puncta that did not colocalize with Lamp1+ lysosomes or bacteria were also apparent (Figure S4C and S4D) and these accumulated at later timepoints (Figure S4E and S4F). To determine if these unassigned APOL3 puncta might represent mitochondria, we undertook kinetic analysis of APOL3 distribution during transient lysosomal damage by pulsing with LLOME, then tracking APOL3^mnGFP^ positioning during the ‘wash-out’ using Lamp1-mCherry and SNAP®-tag-OMP25 (hereafter termed OMP25^646^) to mark lysosomes and mitochondria respectively. After initially forming puncta at lysosomes, APOL3^mnGFP^ foci were predominantly localized to mitochondria when imaged later, 2 hr after removal of LLOME insult (Figure 2D). This corresponded to 5-20% of the total cellular volume of mitochondria being targeted by APOL3 after lysosomal damage (Figure S4G). Imaging at intermediate time points captured the transitionary period, with APOL3 foci being roughly equally distributed between lysosomes and mitochondria (Figure 2D). APOL3-GFP expressed in THP-1 macrophages also trafficked to mitochondria when fed SiO_2_ particles (Figure 2E). Moreover, similar mitochondria-APOL3 structures were apparent after experiencing infectious lysosomal damage induced by cytosol invasion by *Stm* (Figure 2F). Thus, APOL3 translocate to mitochondria and enhance mtDNA release downstream of diverse triggers of sterile or infectious lysosomal damage.

### BAK/BAX-driven MOMP targets APOL3 to mitochondria and is required for mtDNA release during lysosomal damage

We next considered the functional relationship between damaged lysosomes and mitochondria. Upon severe damage, released lysosomal cathepsins promotes the oligomerization of the proapoptotic proteins BAX and BAK on the outer mitochondrial membrane (OMM) to initiate mitochondrial outer membrane permeabilization (MOMP) and intrinsic apoptosis^3,4,23–27^. However, MOMP does not always lead to apoptosis, as sub-lethal MOMP can occur in a minority of mitochondria (termed miMOMP) without detectable cell death^30,46,47^. To determine if transient lysosomal damage triggered MOMP, we used a fluorescent reporter system where co-localization of cytosolic GFP (cytoGFP) and a mitochondrial mCherry (mitoCherry) in the presence of heterodimerizer is used as a readout of MOMP^30^. We found that a 2 hr pulse of LLOME induced miMOMP in approximately 12.5% of mitochondria, which contrasts with ABT-737/Q which induced widespread MOMP (Figure 3A). We then engineered HeLa cells that lack both BAK and BAX and found this abolished the ability of LLOME to elicit miMOMP (Figure 3B). Thus, transient lysosomal damage triggers BAK/BAX-dependent miMOMP in living cells.

**Figure 3.**
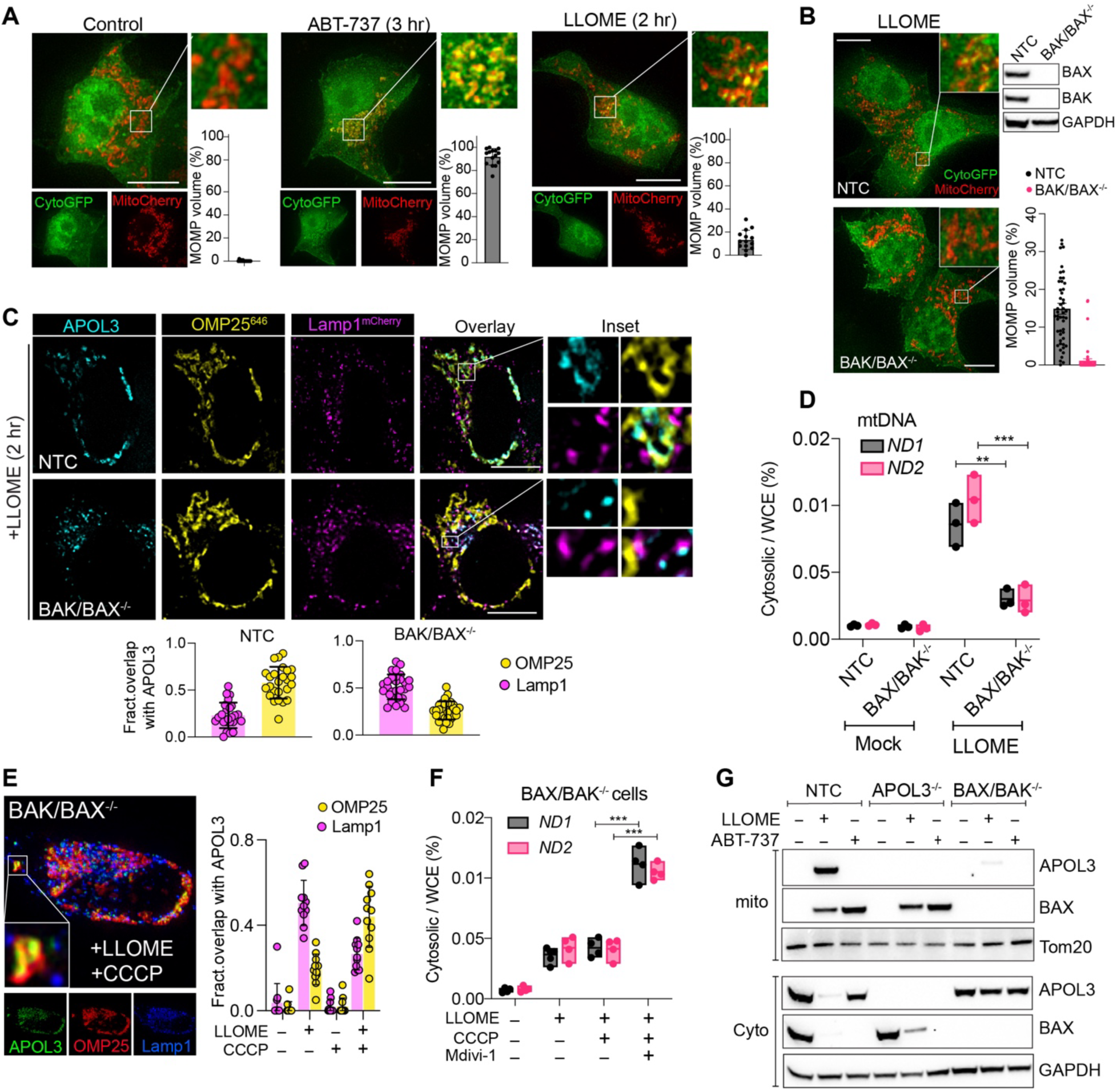
BAK/BAX-induced MOMP is required for APOL3 mitochondrial targeting and mtDNA release. (A, B) IFN-γ-primed HeLa cells expressing cytosolic GFP (CytoGFP) and mitochondrial mCherry (MitoCherry) were treated with LLOME or ABT-737/Q with heterodimerizer and imaged. Total mitochondrial volume (MitoCherry) overlapping with cytoGFP is shown (mean ± SD from 15 (A) or 50 (B) cells. (C) IFN-γ-primed NTC or BAK/BAX^-/-^ cells expressing APOL3-GFP, OMP25^646^, or Lamp1-mCherry treated with LLOME. Fractional overlap for APOL3 is shown (n = 25 cells, error bars ± SD). (D) qPCR of cytosolic mtDNA relative to WCE in IFN-γ-primed NTC or BAK/BAX^-/-^ cells treated with LLOME. (E) IFN-γ-primed BAK/BAX^-/-^ cells as in (C) were treated with LLOME with or without CCCP. Fractional overlap for APOL3 is shown (n = 11 cells, error bars ± SD). (F) qPCR quantification of mtDNA in the cytosol relative to WCE of IFN-γ-primed BAK/BAX^-/-^ cells with the indicated agonist. (G) Representative immunoblots of cytosolic (cyto) or mitochondrial (mito) fractions prepared from IFN-γ-primed HeLa genotypes treated with LLOME or ABT-737/Q. (A), (B), (C), (E) are representative or >3 independent experiments. (D) and (F) are floating bars (mean, min/max, n = 3 or 4 independent experiments). ** P < 0.01; *** P < 0.001 determined by one-way ANOVA. Images are maximum intensity projections (A, B, E) or single *z* planes (C) of Thunder deconvolved widefield images. Scale bar, 5 μm.

This result allowed us to ask if LLOME-triggered MOMP was required for APOL3 mitochondrial targeting. In IFN-γ-primed BAK/BAX^-/-^ cells, APOL3 puncta remained associated with Lamp1^+^ structures and failed to target mitochondria as scene in NTC cells (Figure 3C). BAK/BAX^-/-^ cells were also defective in releasing mtDNA (Figure 3D), activating IRF3 (Figure S5A) and upregulating type I IFN-responsive antiviral ISGs (Figure S5B) following LLOME treatment. Strikingly, treating BAK/BAX^-/-^ cells with LLOME and CCCP (which induces broad mitochondrial damage) mostly rescued APOL3 mitochondrial targeting (Figure 3E) and mtDNA efflux (Figure 3F). This latter effect required mitophagy inhibition with Mdivi-1, consistent with CCCP being an inducer of mitophagy which curtails mtDNA release ^48^. In contrast, inducing MOMP alone with ABT-737 or CCCP in the absence of lysosomal damage did not induce APOL3 puncta formation nor its mobilization to mitochondria (Figure S5C). Cellular fractionation corroborated our microscopy results, showing APOL3 translocation to the mitochondrial fraction during lysosomal damage in a BAX/BAK-dependent manner, but not during mitochondrial damage alone (Figure 3G). In contrast, BAX was mobilized to mitochondria by ABT-737 and by LLOME (although to a lesser degree) in a manner independent of APOL3 (Figure 3G). Together, these data argue that APOL3 is recruited to mitochondria to enhance mtDNA release under specific conditions wherein lysosomal and mitochondrial damage occur in concert.

Because APOL3 has been shown to regulate mitophagy in podocytes^49^, we analyzed whether mitophagy was involved in APOL3 mediated mtDNA release. Using the mitophagy reporter Cox8-mCherry/GFP^50^, we found that either CCCP or LLOME induced mitophagy at similar frequencies in NTC and APOL3^-/-^ cells (Figure S5D). Moreover, APOL3 exhibited enhanced trafficking to mitochondria in mitophagy deficient Parkin (PRKN)^-/-^ cells (Figure S5E). This could be explained by the ability of Parkin to inhibit the pro-apoptotic functions of BAK/BAX^51^, or by shuttling of mitochondria inside autophagosomes before they can be recognized by APOL3^52^. In accordance with PRKN attenuating the APOL3 response, inducing mitophagy with urolithin A concurrent with LLOME suppressed APOL3-dependent IRF3-luciferase activation (Figure S5F). Thus, APOL3 targets mitochondria and promotes mtDNA efflux in a manner that is suppressed by mitophagy.

### APOL3 disrupts the IMM of OMM damaged mitochondria to enhance mtDNA release *in vitro*

APOL3 exerts detergent-like permeabilizing activity against the inner membrane of Gram-negative bacteria^36^. Here, selectivity is enforced through preferential activity against negatively charged, cholesterol-free lipid bilayers– both traits largely specific to microbial membrane^36^. However, inner mitochondrial membranes (IMMs) also retain these same bacterial-like traits due to their endosymbiotic origin^53^. This prompted us to consider if APOL3 exhibits similar permeabilizing activity against the IMM. We generated calcein-loaded liposomes mimicking the lipid composition of either the outer (OMM) - or inner (IMM) mitochondrial membrane. We then added recombinant APOL3 (rAPOL3) and measured permeabilization in real time by detecting the release and de-quenching of calcein. This revealed that liposomes mimicking the IMM but not OMM were rapidly permeabilized by rAPOL3 (Figure 4A). IMM perforation required detergent-like activity because mutating four Phe residues on the hydrophobic face of amphipathic helices 2 and 3 to Ser (denoted APOL3^4F-S^) which disrupts bacteriolysis^36^, abolished calcein release (Figure 4A). The bacterial anionic lipid cardiolipin (CL) is the most abundant IMM-specific lipid relative to the OMM^54^. We titrated CL into PC-scaffold liposomes and exposed them to a gradient of rAPOL3. This revealed that CL content governs the sensitivity of lipid bilayers to rAPOL3 permeabilization (Figure 4B) and binding measured by liposome pull-down assays (Figure 4C). Thus, rAPOL3 selectively permeabilizes IMM-mimetic liposomes through its affinity for CL.

**Figure 4.**
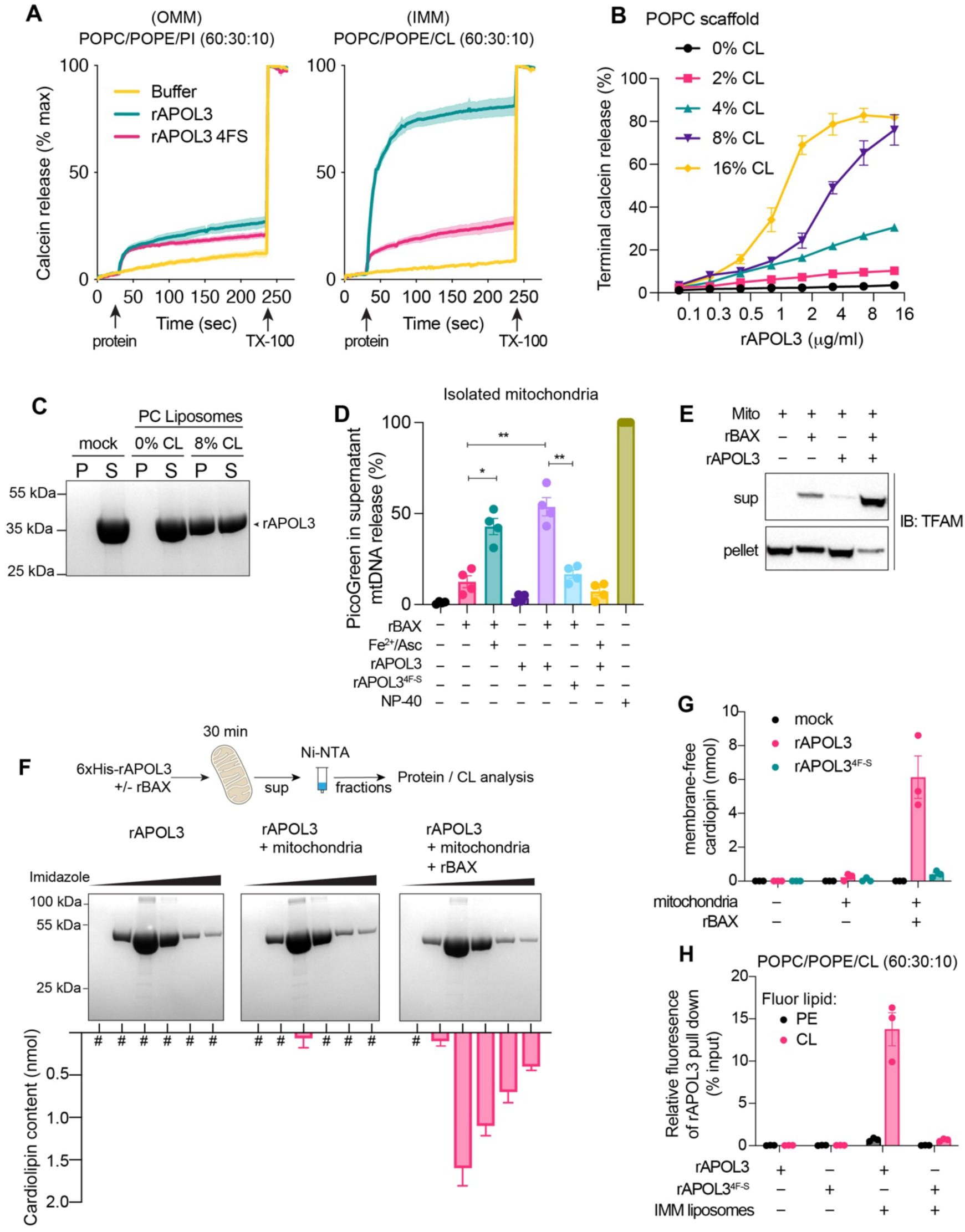
rAPOL3 permeabilizes the IMM of OMM-damaged mitochondria by solubilizing cardiolipin. (A) Time course of calcein leakage from OMM- or IMM-mimetic liposomes treated with 500 nM rAPOL3. (B, C) POPC liposomes with the indicated amount of cardiolipin (CL) treated with a gradient of rAPOL3 for 5 min before terminal calcein release measured (B) and pellets (P) and supernatants (S) stained by Coomassie blue (C). (D, E, F, G) Purified mitochondria treated with rAPOL3 (WT or 4F-S mutant) and/or rBAX for 30 min. mtDNA was release detected by PicoGreen fluorescence in the supernatant (D). Immunoblot for TFAM (mitochondrial transcription factor A) in supernatants or pellets is shown in (E). rAPOL3 was recovered from supernatants by Ni-NTA purification and eluate fractions analyzed by Coomassie blue or CL content (F). Total CL released into supernatant is shown in (G). (H) IMM-mimetic liposomes with fluorescent CL or PE (10% of total lipid) were incubated with rAPOL3 for 30 min and fluorescence of recovered rAPOL3 lipoprotein complexes measured. Data normalized to total fluorescence (input) before addition of protein. (A, B) represent the mean ± SD (n = 3) and are representative of 3 independent experiments. (D, G, H, F) are mean ± SEM from 3 or 4 replicates. Immunoblots and gels are representative of 2 (F) or 3 (C, E) independent experiments. *P < 0.05; ** P < 0.01; *** P < 0.001 determined by one-way ANOVA.

To determine if rAPOL3 can directly permeabilize the IMM of native mitochondria to enhance mtDNA release, we developed a reconstitution system consisting of purified mitochondria that we exposed to recombinant BAX (rBAX), which is sufficient to trigger MOMP *in vitro*^55^, and/or rAPOL3 and measured mtDNA release by PicoGreen^TM^ fluoresence in the supernatant. Adding rBAX alone to mitochondria released 11.2 ± 4.3% of mtDNA; However, supplementing this reaction with rAPOL3 resulted in the majority of mtDNA being released into the supernatant (53.4 ± 8.1%) (Figure 4D). rAPOL3 had minimal effect on mtDNA efflux in the absence of rBAX, in agreement with its inability to permeabilize OMM-mimetic liposomes (Figure 4D). mtDNA release was lost for rAPOL3^4F-S^, and could be phenocopied by inducing oxidative stress using a ROS-generating system Fe(II)SO_4_/ascorbate (Fe^2+^/Asc) which destabilizes the IMM^56,57^ (Figure 4D). Immunoblotting the supernatants and pellets of these reactions found that the release of the mtDNA-binding protein TFAM (transcription factor A, mitochondrial) which packages mtDNA into nucleoids largely phenocopied mtDNA release – with rAPOL3 and rBAX synergizing for maximal TFAM efflux (Figure 4E). Thus, rAPOL3 assists in perforating the IMM to enhance mtDNA efflux from mitochondria whose OMM is already permeabilized by BAX.

### APOL3 destabilizes the IMM by solubilizing cardiolipin

The detergent-like activity of rAPOL3 stems from an ability extract membrane-embedded anionic lipids into soluble lipoproteins particles^36^. To determine if APOL3-CL lipoprotein complexes are formed during permeabilization of the IMM, we used immobilized metal affinity chromatography to recover soluble rAPOL3 from the supernatants of completed mitochondrial reactions that had been incubated with 6xHis-rAPOL3 and rBAX. After extensive washing, we collected eluate fractions and used a fluorometric assay to measure CL that co-purified with rAPOL3. When recovered from the supernatants of permeabilized mitochondria, rAPOL3 positive fractions contained abundant CL that was lost if either mitochondria or rBAX were omitted from the reaction (Figure 4F). Size exclusion chromatography revealed most of the CL released from permeabilized mitochondria was in fact contained within rAPOL3 lipoprotein complexes which were considerably larger in size than input lipid-free rAPOL3 (Figure S6A and S6B). Mutating the active site of rAPOL3 to APOL3^4F-S^ negated the release of membrane-free CL (Figure 4G). To determine if if APOL3 selectively extracts CL from the IMM, we spiked IMM liposomes with fluorescent (TopFluor®) CL or PE to 10% of total lipid while preserving the lipid ratio. We then measured how much of each fluorescent lipid was contained within rAPOL3 lipoprotein complexes recovered from permeabilized mitochondrial. rAPOL3 solubilized 13.7 ± 4.8% of the input CL, but less than 1% of input PE that composed the original liposome bilayer (Figure. 4H). These data argue that APOL3 directly permeabilizes the IMM by solubilizing CL.

### Detergent-like activity and mitochondrial localization are required for APOL3 to enhance mtDNA release inside cells

We enlisted biochemically defined mutants to discern which APOL3 functions are required to release mtDNA inside cells. We rescued APOL3^-/-^ cells with WT or APOL3 in which Leu137-Ala138-Pro-139 is mutated to Gln137-Ser138-Ser138 (APOL3^GSS^) – which mislocalizes APOL3^GSS^ to the cytosol during lysosomal damage^36^. APOL3^GSS^ failed to form puncta on mitochondria (Figure 5A) and failed to rescue IFN-γ-enhanced mtDNA release (Figure 5B) or IRF3 activation (Figure 5C) release upon LLOME treatment. Next, we rescued APOL3^-/-^ cells with WT or APOL3^4F-S^, which is defective in permeabilizing IMM liposomes and native mitochondria (Figure 4A and 4C). After inducing lysosomal damage, APOL3^4F-S^ still formed puncta at mitochondria (Figure 5A) but failed to rescue both LLOME-triggered mtDNA release (Figure 5B) and IRF3 activation (Figure 5C). We then asked if similar APOL3 functions were required for the response infectious lysosomal damages. WT but not APOL3^4FS^, or APOL3^GSS^ rescued ISG induction triggered by infecting IFN-γ-primed APOL3 cells^-/-^ cells with cytosol-invasive *Stm* or AdV (Figure 5D). THP-1 macrophages also required active and mitochondrial localized APOL3 to efficiently release mtDNA and produce IFN-β when fed SiO_2_ particles (Figure 5E and 5F). Thus, both APOL3 detergent-like activity and its location at mitochondria is required to promote mtDNA release inside cells.

**Figure 5.**
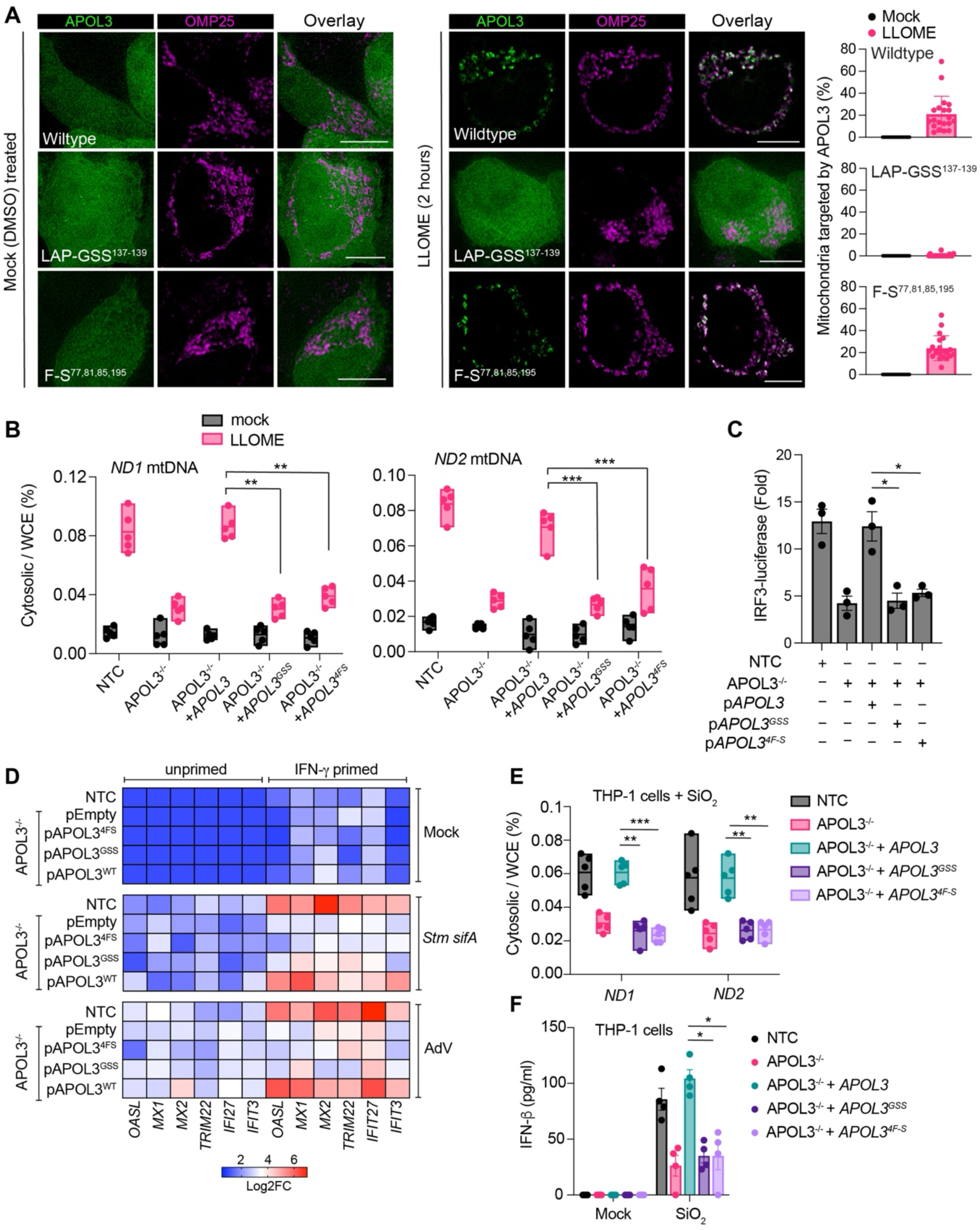
Detergent-like activity and mitochondrial localization are required for APOL3 to enhance mtDNA release. (A) IFN-γ-primed HeLa cells co-expressing the indicated variant of APOL3-GFP and OMP25^JF646^ treated with LLOME. Volume of OMP25 overlapping with APOL3 is shown (n = 20 cells, mean ± SD). Images are single *z* planets of Thunder deconvolved widefield images and are representative of 3 independent experiments. Scale bar, 5 nm. (B and C) IFN-γ-primed APOL3^-/-^ cells rescued with the indicated APOL3 variant were treated with LLOME (2 hr) and mtDNA in the cytosol relative to WCE determined by qPCR (B) or luminescence detected in cells expressing IRF3-luciferase after an additional 4 hr (C). (D) APOL3^-/-^ or NTC cells +/- IFN-γ priming rescued with the indicated *APOL3* variant and infected with *Stm* Δ*sifA* for 6 hr or AdV for 12 hr. FC relative to mock treated for each gene was determined by qPCR (n = 2). (E and F) IFN-γ/IL-1β-primed APOL3^-/-^ THP-1 macrophages rescued with the indicated *APOL3* variant were fed SiO_2_ particles to induce phagosomal rupture. Cytosolic mtDNA relative to WCE was determined by qPCR (E), or IFN-β measured by ELISA after 24 hr (F). (B) and (E) are floating bars (mean, min/max, n = 5 independent experiments). (C) and (F) are mean ± SEM from 3 or 4 independent experiments. *P < 0.05; ** P < 0.01; *** P < 0.001 determined by one-way ANOVA.

### APOL3 neutralizes an IMM barrier that impedes mtDNA release during MOMP

MitoDAMPs are spatially separated within mitochondria – cytochrome *c* is loosely tethered within the intermembrane space, whereas mtDNA is fully enclosed by the IMM^29^. BAX/BAK macropores on the OMM mediate rapid and complete release of cytochrome *c*^58^, but can also allow the IMM to herniate into the cytosol and release mtDNA^45,59^. However, herniation and permeabilization of the IMM is a relatively rare event that occurs after cytochrome *c* release^45^. This suggests that the IMM stays sufficiently intact during MOMP to impede wholesale mtDNA release until after apoptotic caspases are activated by cytochrome *c*, rendering apoptosis immunogenically silent^43^. Indeed, MOMP typically results in a type I IFN response only under caspase-inhibited conditions^43,45,59^. We hypothesized that by selectively permeabilizing the IMM, APOL3 helps negate the natural bias of mitochondria to release cytochrome *c* more efficiently than mtDNA, thereby enabling miMOMP to release sufficient mtDNA to activate cGAS. Consistent with our sub-lethal treatment regiment, we detected only a slight increase in Annexin-V staining (an indicator of apoptosis) upon LLOME exposure that was partially blocked by the pan-caspase inhibitor QVD-OPh (Figure 6A). This contrasts with activating BAK/BAX directly with ABT-737 which induced robust apoptosis that was completely blocked by pan-caspase inhibition (Figure 6A). Caspase inhibition did not significantly impact APOL3-mediated IFN-β production triggered by lysosomal damage, but dramatically increased IFN-β release during BAK/BAX-mediated apoptosis (Figure 6B). This indicates the APOL3-mediated IFN-β response occurs without robust caspase-dependent apoptosis.

**Figure 6.**
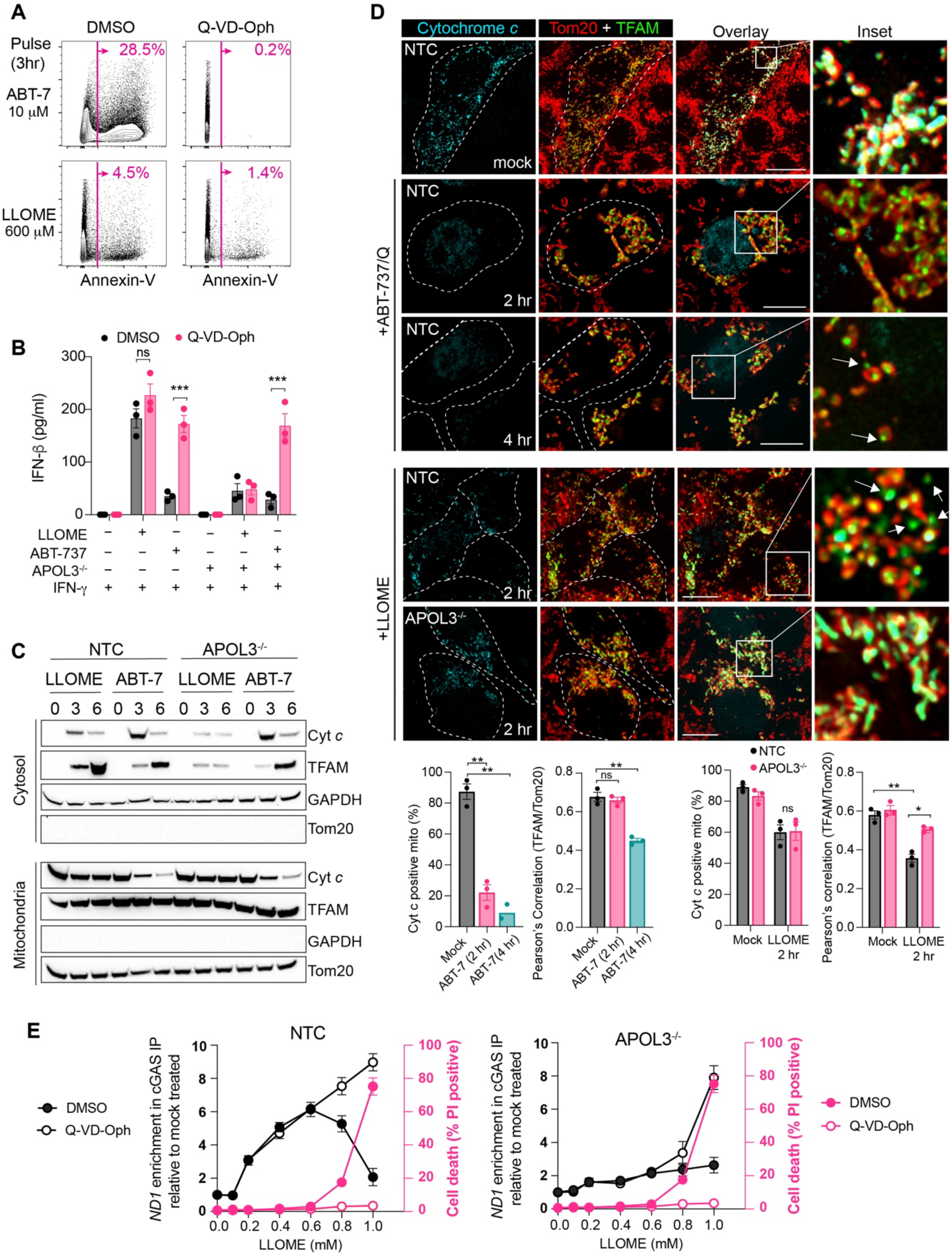
APOL3 neutralizes an IMM barrier that normally impedes mtDNA release during MOMP. (A, B) IFN-γ-primed HeLa cells pulsed with ABT-737 or LLOME with or without pan-caspase inhibition (Q-VD-Oph) and analyzed by Flow cytometry for Annexin-V staining or IFN-β production by ELISA, both after 24 hr. (C) Time course immunoblots of cytosolic or mitochondrial fractions from IFN-γ-primed HeLa cells incubated with LLOME or ABT-737. Blots are representative of 3 independent experiments. (D) IFN-γ-primed NTC or APOL3^-/-^ cells expressing cytochrome *c* – GFP and TFAM-mScarlet treated with ABT-737/Q or LLOME. Images are maximum intensity projections, representative of 3 independent experiments. Quantification is mean ± SEM from 3 independent experiments (n = 20 cells per replicate). Scale bar, 5 μm. (E) IFN-γ-primed NTC or APOL3^-/-^ HeLa cells expressing FLAG-cGAS were pulsed with LLOME (2 hr) at the indicated concentration. 2 hr later cell death was measured by PI uptake (right y axis, rose) followed by qPCR for mtDNA in FLAG immunoprecipitants (left y axis, black). Where indicated treatments included Q-VD-Oph. (B) and (E) are mean ± SEM from 3 independent experiments. *P < 0.05; ** P < 0.01; *** P < 0.001 determined by one-way ANOVA.

Next, we conducted kinetic analysis of mitochondrial DAMP release into the cytosol by cellular fractionating and immunoblotting. When compared to inducing apoptosis, LLOME effluxed less cytochrome *c* but more TFAM into the cytosol, and this was largely dependent on APOL3 (Figure 6C). We then undertook temporal microscopy analysis of mitochondrial integrity in IFN-γ-activated cells expressing cytochrome *c*-GFP and TFAM-mCherry. In agreement with other reports^45,59,60^, ABT-737/Q induced rapid cytochrome *c*-GFP loss from mitochondria that was mostly complete by 2 hr, a timepoint where TFAM nucleoids were still contained within Tom20 stained OMM (Figure 6D). Cytosolic TFAM and IMM herneation became apparent later, 4 hr after inducing apoptosis (Figure 6D). In contrast, LLOME triggered concurrent cytochrome *c* and TFAM nucleoids efflux, as both were released from a subset of mitochondria and present in the cytosol by 2 hr (Figure 6D). Critically, deletion of APOL3 suppressed the efflux of TFAM without significantly effecting cytochrome *c* release (Figure 6D).

Based on these results, we predicted that a reliance on APOL3 could be used to uncouple mtDNA release that occurred in living cells from that which occurs in dying cells. To test this, we measured the amount of mtDNA bound to FLAG-cGAS at various LLOME doses in both NTC and APOL3^-/-^ cells with or without pan-caspase inhibition and simultaneously measured cell death by propidium iodide (PI) staining. This revealed an APOL3-dependent and -independent effect that segregated based on the degree of lysosomal injury. At low LLOME doses where no cell death occurred, cGAS bound mtDNA in NTC cells but much less so in APOL3^-/-^ cells (Figure 6E). Whereas at high doses of LLOME where significant cell death ensued, cGAS bound mtDNA equally well in both NTC and APOL3^-/-^ cells, but only under caspase-inhibited conditions (Figure 6E). Together, these data suggest that when activated by lysosomal damage, APOL3 lowers the volume of MOMP needed to induce a CGAS-mtDNA interaction to sub-lethal levels. Accordingly, In APOL3^-/-^ cells, the amount of lysosomal damage that induced an equivalent mtDNA-cGAS interaction also triggered apoptosis which suppressed cGAS activation (Figure 6E).

### Cardiolipin externalization recruits APOL3 from lysosomes to mitochondria

We next investigated the mechanism whereby APOL3 traffics to mitochondria. Mitochondria that are permeabilized by BAX/BAK externalize cardiolipin from the IMM to the OMM via the phospholipid scramblase-3 (PLSCR3) ^61,62^. Considering the affinity of rAPOL3 for CL *in vitro*, we asked if lysosomal damage triggered CL externalization on the OMM, and if this served as a signal for recruiting APOL3. We stained for cardiolipin in OMP25^646^ expressing cells using differential saponin permeabilization. Because saponin acts by forming complexes with cholesterol, cholesterol-rich membranes like the plasma membrane are efficiently permeabilized whereas cholesterol-poor membranes like mitochondria remain impenetrable^63^. Saponin-enabled signal thus serves as a readout of OMM-exposed cardiolipin. Using this method, we found that LLOME-triggered positive cardiolipin staining on mitochondria (Figure 7A) that could be negated by genetic deletion of PLSCR3 (Figure 7B). We then conducted Airyscan super-resolution imaging using knock-in APOL3^HA^ cells pulsed with LLOME and identified saponin-exposed cardiolipin on mitochondrial membranes that overlapped with regions of high APOL3 intensity (Figure 7C). Next, we asked whether CL externalization was required for APOL3 mitochondrial targeting. In PLSCR3^-/-^ cells pulsed with LLOME, APOL3-GFP puncta remained associated with Lamp1^+^ structures and failed to target mitochondria (Figure 7D). Accordingly, PLSCR3^-/-^ cells exhibited reduced cytosolic mtDNA (Figure 7E), IRF3 activation (Figure S7A) and ISG upregulation (Figure S7B) when treated with LLOME. PLSCR3^-/-^ THP-1 macrophages were also defective in producing IFN-β but not IL-1β when fed SiO2 particles (Figure S7C). We next isolated mitochondria from NTC or PLSCR3^-/-^ cells and directly assayed their sensitivity to rAPOL3 in the presence of rBAX. Mitochondria from PLSCR3^-/-^ cells were on average 6.2-fold more resistant to APOL3-mediated permeabilization (LD_50_ = 8.7 μM) than mitochondria harvested from NTC cells (LD_50_ = 1.4 μM) (Figure 7F). Collectively, these data show that lysosomal damage induces PLSCR3-dependent CL externalization which recruits APOL3 to promote mtDNA release.

**Figure 7.**
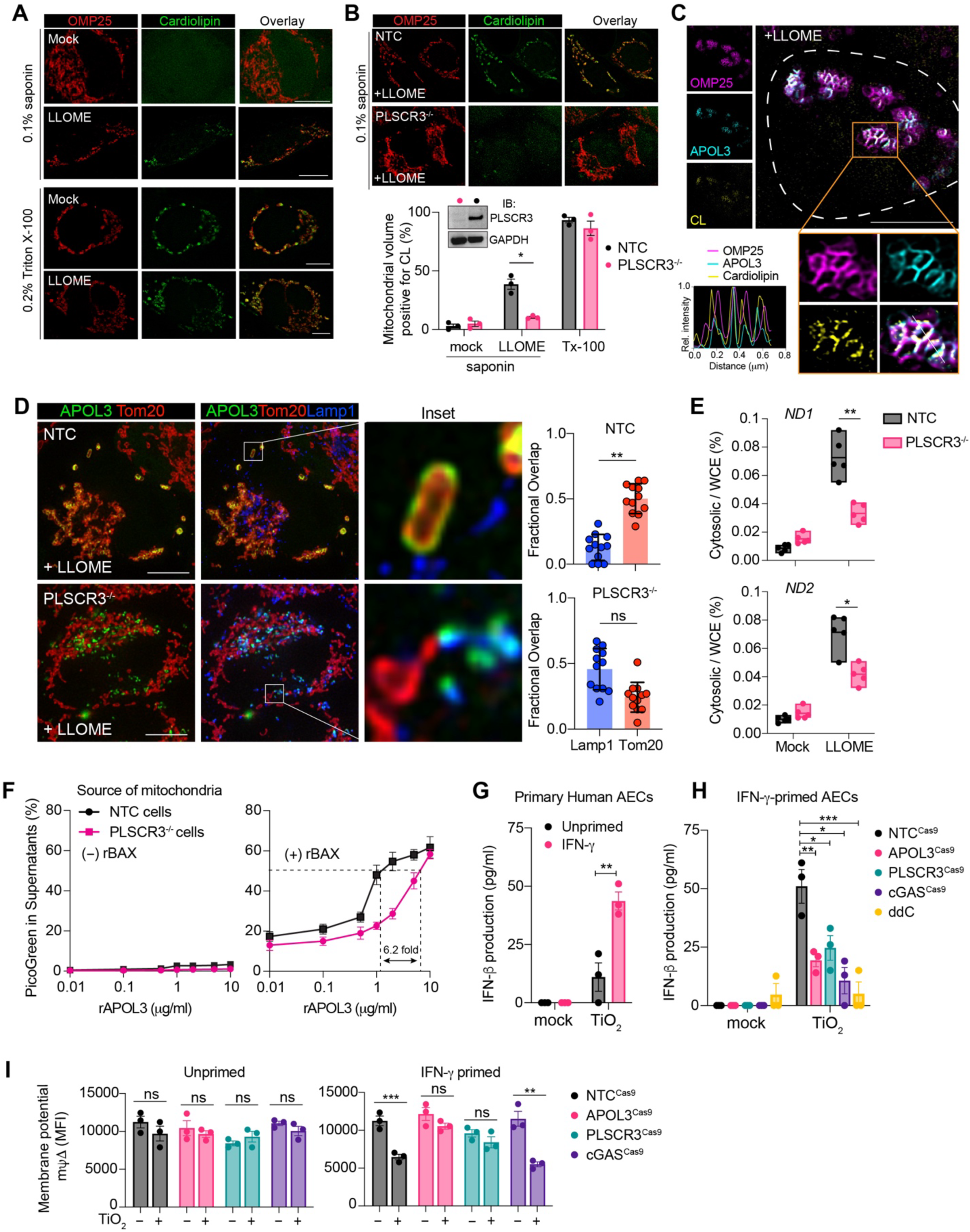
Cardiolipin externalization recruits APOL3 from lysosomes to mitochondria. (A and B) Cardiolipin immunofluorescence (single *z* planes) in IFN-γ-primed wildtype HeLa cells (+/- LLOME) permeabilized with saponin or Trition-X100 (A) or in NTC and PLSCR3^-/-^ cells + LLOME using saponin permeabilization (B). Quantification represents mean ± SEM from 3 independent experiments (n = 20 cells per replicate). Immunoblots (IB) are representative of 2 experiments. (C) Representative Airyscan image of IFN-γ-primed LLOME-treated APOL3^mnGFP^ cells stained for OMP25 with JF646 and CL. Single *z* slices and line profile is shown. (D) APOL3-GFP expressed in IFN-γ-primed NTC or PLSCR3^-/-^ cells treated with LLOME and immunostained for Tom20 and Lamp1. Fractional overlap for APOL3 is shown (n = 12 cells, representative of 3 independent experiments, error bars ± SD). Maximum intensity projections of Thunder deconvolved *z* stacks are shown. (E) qPCR quantification of mtNDA in the cytosol versus WCE in IFN-γ-primed HeLa genotypes treated with LLOME. Shown are floating bars (mean, min/max, n = 5 independent experiments). (F) Mitochondria isolated from NTC or PLSCR3^-/-^ cells treated with rAPOL3 +/- rBAX and mtDNA release detected by PicoGreen fluorescence. Data are mean ± SEM (n = 3). (G, H, I) Primary human airway epithelial cells (AECs) primed with IFN-γ (or not) were treated with TiO2 nanoparticles (24 hr) and IFN-β measured in the supernatants by ELISA (G). AECs were transduced with lentivirus encoding Cas9 and sgRNAs targeting *APOL3*, *PLSCR3*, or *cGAS* or cultured with ddC prior to treatment (H). Alternatively, AECs were stained with Tetramethylrhodamine Methyl Ester (TMRM) after 12 hr to probe mitochondrial membrane potential (ΔΨm) and quantified by flow cytometry (MFI: median fluorescence intensity). (G, H, I) represent the mean ± SEM with AECs prepared from three different organ donors. * P < 0.05; ** P < 0.01; *** P < 0.001 determined by one-way ANOVA (B, E, G, H, I) or two-tailed unpaired *t* test (D).

Lastly, we assessed whether the PLSCR3-APOL3-cGAS pathway was activated in a human primary cell model of lysosomal damage. Airway epithelial cells (AECs) are continually exposed to inhaled particulates which can induce lysosomal damage and dysfunction leading to lung injury. We isolated primary AECs from human organ donors and assessed their response to ultrafine TiO_2_ nanoparticles – a main component of air pollution that triggers inflammatory cytokines^64,65^. TiO2-treated AECs from three different donors produced significantly more IFN-β when first primed with IFN-γ (Figure 7G). This heightened response was suppressed by lentiviral CRISPR/Cas9 knockdown of PLSCR3, APOL3, or cGAS and was blocked by mtDNA depletion with ddC (Figure 7H). We then analyzed mitochondrial health by detecting the electrical potential maintained across the IMM with Tetramethylrhodamine, methyl ester (TMRM) which accumulates within mitochondria with intact membrane potential. TiO2 particles reduced mitochondrial membrane potential specifically in IFN-γ primed AECs in a manner dependent on APOL3 and PLSCR3 but not cGAS (Figure 7I). These data support our model that PLSCR3 and APOL3 are upstream effectors of mitochondrial damage, whereas cGAS is the downstream detector. Moreover, they highlight how inflammatory signals like IFN-γ elicit innate immune effectors that pre-condition human tissues to respond with more urgency to subsequent injury or infection.

## DISCUSSION

In this study, we describe how human cells deploy the antibacterial protein APOL3 against an endogenous host organelle to promote the selective breakdown of mitochondrial endosymbiosis. This type of ‘introspective’ function for APOL3 was triggered specifically by lysosomal damage and potentiated a type I IFN response to both injury and infection. Transient lysosomal damage activates sub-lethal apoptotic MOMP via BAK/BAX macropores; however, the IMM normally stays sufficiently intact in these mitochondria to prevent wholesale mtDNA release and limit type I IFN production. We found that IFN-γ priming induced the expression of APOL3 which recognizes externalized cardiolipin on these damaged mitochondria undergoing MOMP. APOL3 selectively permeabilizes the IMM by solubilizing cardiolipin to enhance mtDNA release and downstream cGAS activation. We showed that by facilitating mtDNA release on a per-mitochondria level, APOL3 lowered the total volume of MOMP needed to trigger cGAS activation to sub-lethal levels – but only when concurrent with lysosomal damage. For resting or IFN-γ-primed APOL3^-/-^ cells, the amount of lysosomal damage that induced an equivalent mtDNA-cGAS interaction was concomitant with apoptosis which inhibited downstream IFN production.

There are several mechanisms that promote mtDNA release, including BAK/BAX macropores^45^, VDAC channels^60,66^, vesicularization^67^, and the pyroptosis effector Gasdermin D^68,69^. APOL3 is unique among these pathways in that it specifically enhances IMM permeability – which our data suggests is rate-limiting for mtDNA efflux in living cells. Reflecting the natural inefficiency of IMM permeabilization during MOMP, studies that have observed mtDNA efflux and cGAS activation in living cells generally use caspase inhibitors to prevent cell death^45,59^, or enlist long-term pathological conditions like cellular senescence ^32,70^, mtDNA replication stress^22,71^, or animal models harboring genetic defects in key metabolic pathways^60,67,71^. In contrast, APOL3 is activated by an endogenous danger signal and appears to play a role in generating a type I IFN response and mobilizing antiviral immunity independently of cell death.

Pathogens of all phylogenic classes cross the endolysosomal barrier when entering their intracellular niche^72,73^. Reacting to a breach of this membrane is therefore a way to trigger IFN production and mobilize host defense independently of pathogen structure. Such a mechanism would be less susceptible to evolutionary escape and be effective against emerging zoonoses that could otherwise go undetected by the host^74,75^. Indeed, genetic and proteomic screens have identified possible roles for APOL3 and related APOL family members to include restriction activity against a remarkably wide array of structurally diverse pathogens, including African trypanosome parasites^76^, *Salmonella* and *Shigella* bacteria^36^, and viruses such as adenovirus, encephalitis virus, parainfluenza virus, coxsackie virus, poliovirus, Sinbus virus, and SARS-CoV-2 ^77–81^. In this context, our results provide a possible explanation for the sweeping anti-pathogen activity of this family in that their function intersects with an endogenous process perturbed by infection (organelle damage) rather than solely targeting the pathogen itself.

As the engines of anabolism and catabolism respectively, lysosomes and mitochondria exhibit extensive crosstalk to regulate cellular homeostasis^82^. This occurs most dramatically during mitophagy where damaged mitochondria are delivered to lysosomes for disposal. However, our data suggests it is unlikely mitophagy plays a major role in delivering APOL3 from damaged lysosomes to mitochondria. High-resolution imaging has demonstrated dynamic inter-organelle membrane contact sites between mitochondria and lysosomes that are distinct from mitophagy^83^. These contacts persist during lysosomal damage and increasing their frequency triggers downstream mitochondrial responses in macrophages^19^. Whether contact sites serve as an inter-organelle conduit for APOL3 remains unclear, but it is notable that 15% of lysosomes are in contact with mitochondria at a given point in time^83^, which correlates with both the frequency of MOMP and mitochondria-localized APOL3 during lysosomal damage. Lysosomes also engulf defective mitochondrial nucleoids from the herniating IMM in a macroautophagy-like process ^21^. Retrograde transfer of APOL3 from lysosomes to mitochondria via this pathway could also explain our results, though time-lapse high-resolution and structural imaging studies would be required to address this.

Our study highlights how the inflammatory environment of the niche dictates how a cell responds to infection and injury. IFN-γ likely represents an alarm signal of increased urgency given that it is made by lymphocytes in a cell-extrinsic manner and typically requires additional signals like antigen or co-stimulation to regulate its synthesis and release^33^. In this regard, that *APOL3* is upregulated by IFN-γ and not IFN-β - the principal output of the pathway under its control – serves to prevent a pathological pro-inflammatory feed-forward loop. As a corollary, our study suggests that by potentiating the cellular responses to lysosomal damage, IFN-γ response signatures that dominate the transcriptomes of many pro-inflammatory or autoimmune diseases may participate in disease pathogenesis rather than simply acting as a marker of disease. Indeed, lysosomal damage does not just occur in response to external insults like infection or environmental particles, but is continually occurring as consequence of endogenous factors including membrane lipid oxidation^84–86^, protein aggregation^19,87,88^, and cellular quiescence^89^. Though most cells typically detect and repair this damage to survive, our study suggests that even transient lysosomal leakage could have pronounced immunogenic effects if it occurs within an IFN-γ pre-conditioned tissue niche.

### Limitations of the study

We found that APOL3 enhances mtDNA release by permeabilizing the IMM. It remains unclear whether APOL3 invades through OMM BAK/BAX macropores to permeabilize the IMM, or whether it exerts its activity after the IMM herniates through the OMM. We did not find conclusive evidence of APOL3 positioned inside OMM markers, so we favor the later explanation. However, more detailed structural analysis via super-resolution microcopy or Cryo-EM studies will be required to confirm this.

## RESOURCE AVAILABILITY

### Lead contact

Further information and requests for resources and reagents should be directed and will be fulfilled by the lead contact, Ryan Gaudet

### Materials availability

All unique/stable reagents generated in this study are available from the lead contact with a completed materials transfer agreement.

### Data and code availability

- RNA sequencing data will be deposited to GEO upon publication
- Original western blot images and microscopy images will be deposited at Mendelay Data upon publication
- This paper does report original code

## ACKNOWLEDGEMENTS

This work was supported by the National Institutes of Health (NIH) grant DP2 AI177904 to R.G.G. and NIH grants AI106697 and AI128949 to D.F. R.G.G was supported by the Kinship Foundation through the Searle Scholar Program. D.A.R and A.O were supported by the Columbia University Graduate Training Program in Microbiology and Immunology (T32 AI106711 to D.A.R) and in Cellular, Molecular and Biomedical Studies (T32 GM145766 to A.O). Research reported in this publication was performed in the CCTI Flow Cytometry Core, supported in part by the Office of the Director, National Institutes of Health under awards S10OD030282. These studies used the Confocal and Specialized Microscopy Shared Resource of the Herbert Irving Comprehensive Cancer Center at Columbia University, funded in part through the NIH/NCI Cancer Center Support Grant P30CA013696. We wish to thank the donor families for their generosity and the exceptional efforts of the transplant coordinators and staff of LiveOnNY.

## AUTHOR CONTRIBUTIONS

The project was conceived and supervised by R.G.G Experiments were designed and performed by R.G.G, D.A.R, A.G, H.S, and J.H. B.V assisted with generating key reagents. D.F procured lung tissues from human organ donors. R.G.G wrote the manuscript with input from all authors.

## METHODS

### KEY RESOURCES TABLE

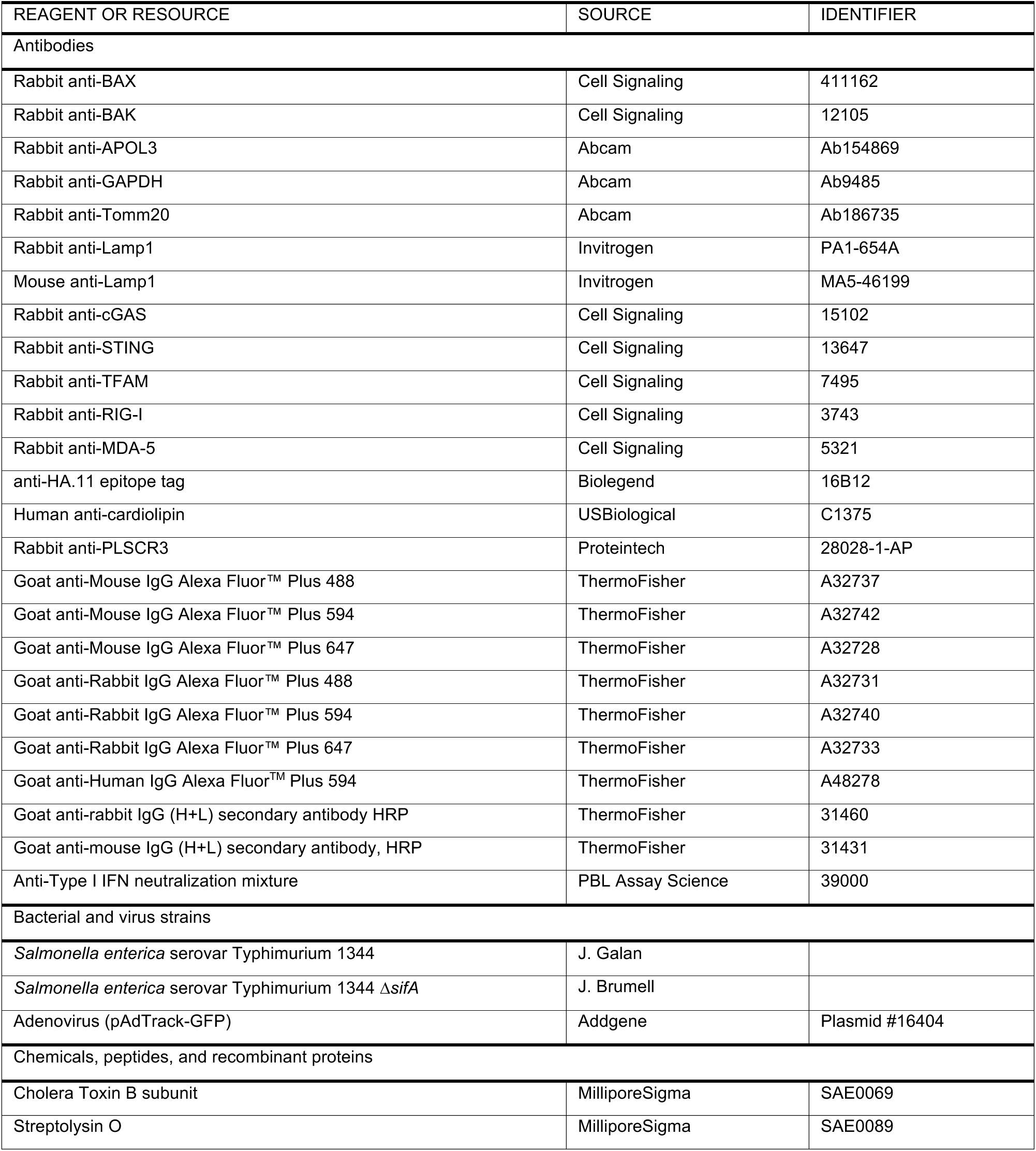
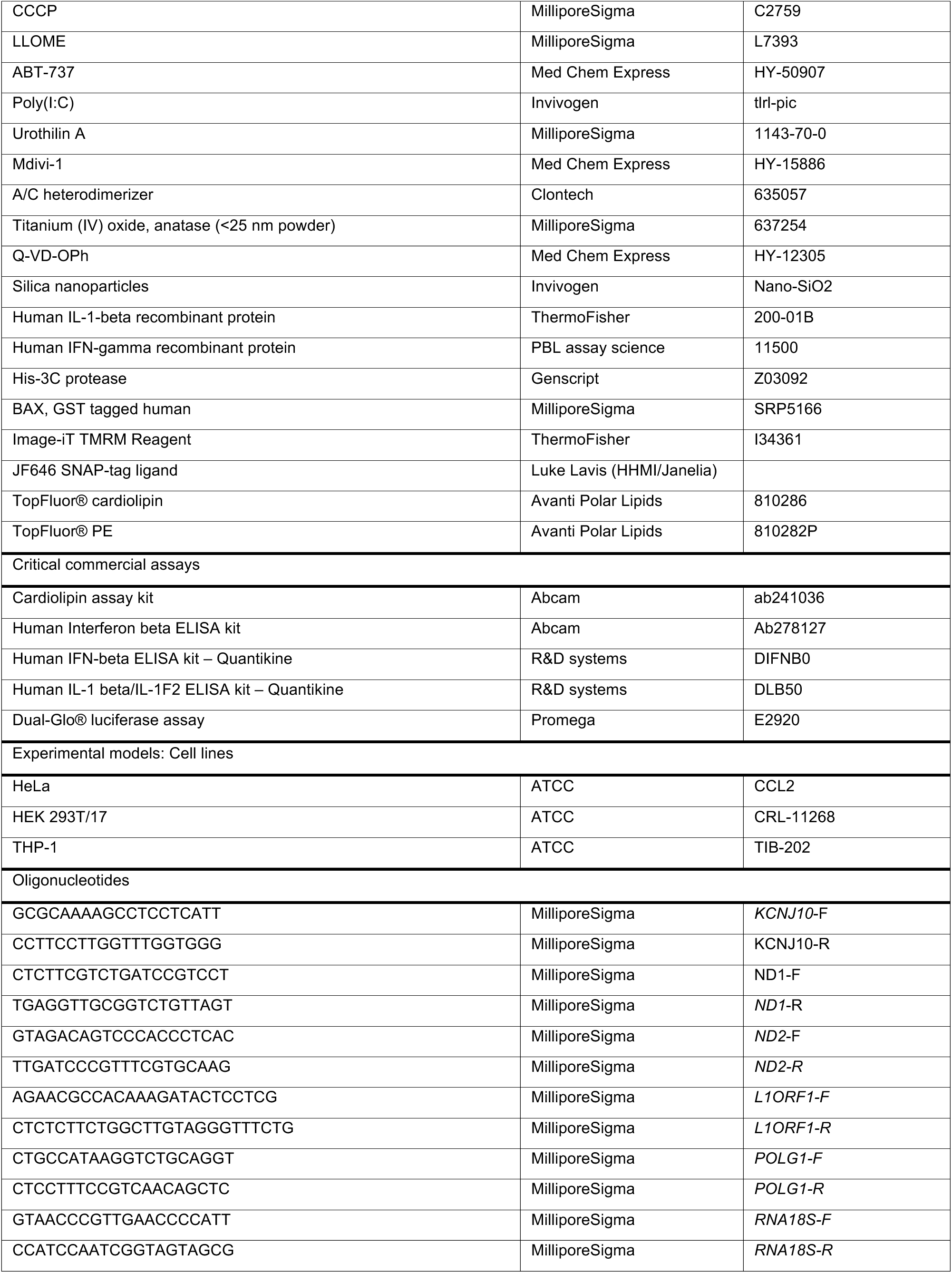
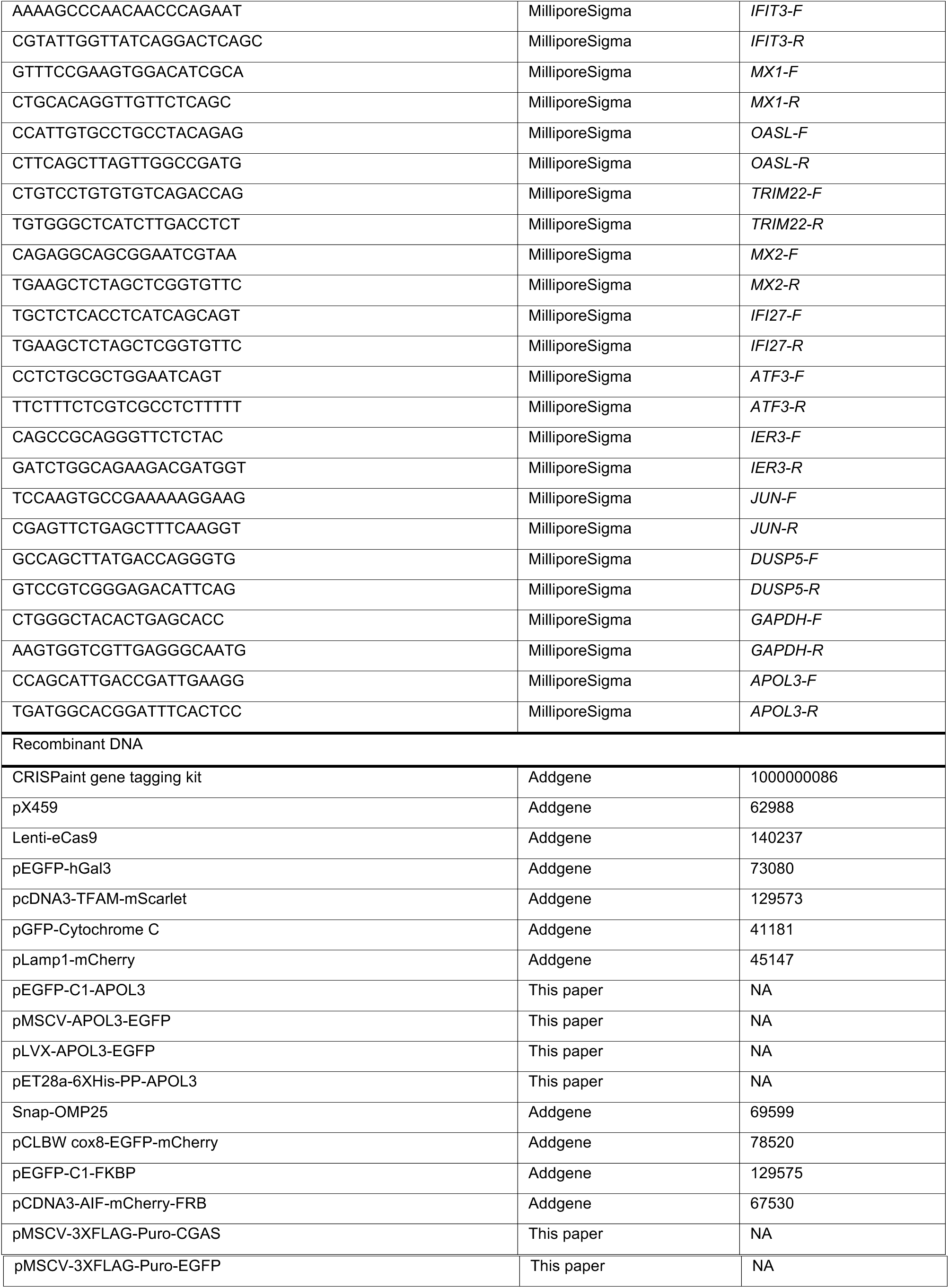

## METHOD DETAILS

### Generation of APOL3 Knock-in cells

We used the CRISPaint^90^ gene tagging kit (Addgene #1000000086) to tag endogenous APOL3 protein with mnGFP or HA. pCRISPaint-mNeon (plasmid #174092) and pCRISPaint-HA (plasmid #67171) were used as donor plasmids. To generate a plasmid expressing sgRNA against human *APOL3*, complementary DNA oligonucleotides (IDT) encoding the crRNA were annealed, phosphorylated and ligated into Esp3I linearized MLM3636 (Addgene #43860), a gift form Keith Joung. pCAS9-mCherry-Frame +2 (plasmid #66941) was used to specify the insertion reading frame. A total of 2 μg of DNA was transfected into HeLa cells in 6 well plates (TransIT-HeLaMONSTER transfection kit; MIRUS) using the following ratio: 1 μg donor plasmid, 0.5 μg frame selector plasmid, and 0.5 μg target selector plasmid. After 48 h, cells were sorted for mCherry positive cells to isolate transfectants. Note that because APOL3 is not constitutively expressed, puromycin selection cannot be used to select for transfected cells. Cells were cloned by single-cell dilution and screened by PCR with a forward primer specific for the 3′ end of the *APOL3* coding sequence, and reverse primers for the 3′ UTR (denote WT) or the HA/mnGFP tag sequence.

### Generation of CRISPR-deleted cell lines

For CRISPR-edited deletions, HeLa or THP-1 cells were transfected with pX459 (Addgene # 62988; a gift from F Zhang) using Mirus LT1 (MIRUS) reagent for HeLa cells or Lipofectamine LTX reagent (ThermoFisher) for THP-1 cells. sgRNA sequences were the following: *APOL3*: 5′-CGC AGT CAC GAA TCT CTT CC-3′, *PLSCR3*: 5′-ACA GGC TAC TTG CCC CCC AA-3′, *PRKN*: 5′-GTG TCA GAA TCG ACC TCC AC-3′, *CGAS*: 5′-*AAG TGC GAC TCC GCG TTC AG*-3′ and 5′-CGC ATC CCT CCG TAC GAG AA-3′, *STING*: 5′-GCG GGC CGA CCG CAT TTGGG -3′ and 5′-CAT ATT ACA TCG GAT ATC TG-3′, *RIGI*: 5′-GGG TCT TCC GGA TAT AAT CC-3′ and 5′-TTG CAG GCT GCG TCG CTG CT-3′, *TLR3*: 5′-TTC GGA GCA TCA GTC GTT GA-3′ and 5′-CAT GCA CTC TGT TTG CGA AG -3′, *MDA5*: 5′-CGA ATT CCC GAG TCC AAC CA-3′ and 5′-GCT TCT AGT TAG AGA CGT CT-3. To generate BAK/BAX double deletions, cells were co-transfected with pX459 harboring sgRNA targeting *BAX* 5′-AGT AGA AAA GGG CGA CAA CC -3′, and *BAK* 5′-GCC ATG CTG GTA GAC GTG TA -3′. Selection was achieved with puromycin (1 μg/ml) for 48 hours. Surviving cells were expanded into media lacking puromycin for 48 hours then subjected to limiting dilution to obtain single colonies where applicable. Colonies were screened first by PCR, then by Western blot, and when appropriate the genotype of each positive clone was determined by Sanger sequencing. For achieving primary cell polyclonal depletions, the same sgRNA sequences were cloned into Lenti-eCas9 (Addgene # 140237, a gift from Urs Greber^91^). Primary airway epithelial cells were transduced with lentivirus at a 1/25 dilution, selected for 48 hr with 2 μg/ml puromycin, then used directly for the experiment.

### RNA sequencing

HeLa NTC or APOL3^-/-^ cells were plated overnight in 6 well plates to reach 70% confluency. Cells were primed (or not) with 500 U/ml IFN-γ for 18 h, after which cells were washed 3X in in PBS and media replaced with fresh DMEM (10% FBS) without cytokines. Selected wells were pulsed with LLOME (600 μM, 2 hr) or DMSO control, rinsed with PBS, then incubated in fresh recovery media without LLOME or cytokines for an additional 8 hr. RNA was then harvested using Quick-RNA^TM^ MiniPrep Plus (Enzo) according to manufacturer instructions. Total RNA was sequenced using PolyA selection for mRNA species with Genwiz® (Azenta Life Sciences). Raw paired-end RNA-seq reads were assessed for quality using FastQC and aligned to the Homo sapiens GRCh38 primary assembly using STAR (version 2.7.11b) with a two-pass mapping strategy. Paired-end reads were processed directly from compressed FASTQ files, and alignments were output as BAM files sorted by genomic coordinates. Gene-level counts were obtained using featureCounts with the same GTF annotation. Differential expression analysis was carried out in Python using PyDESeq2 (version 0.3.5). Raw counts were normalized via the median-of-ratios method and differential expression was assessed using Wald tests, with p-values adjusted by the Benjamini–Hochberg procedure. Volcano plots were generated using Python’s matplotlib and seaborn libraries to visualize differential expression data, with thresholds for significance indicated on the plots.

### Isolation and fractionation of DNA for analysis by qPCR

mtDNA was isolated from whole cell or cytosolic fractions and quantified by qPCR as described previously^92^. Briefly, 10 cm dishes were treated with the indicated ligands, trypsinized and divided into two tubes for the isolation of DNA from whole cell extract and cytosol respectively. For whole cell extract, pellets were lysed in 10 mM Tris pH 8.0 containing 1% SDS at 95 °C for 5 min. To isolate the cytosol, cell pellets were resuspended in Buffer A (20 mM HEPES pH 7.4/KOH, 200 mM sucrose, 50 mM KCl, 10 mM NaCl, 1 mM EGTA, 2 mM MgCl_2_, 1X Halt^TM^ protease inhibitor cocktail) containing 20 μg/ml digitonin and incubated at 4 °C for 10 min on a tube rotator. Supernatant was removed, then spun at max speed for 15 min to pellet any remaining organelles and stored as cytosol extract. For some experiments, the organelle pellet was washed twice with Buffer A, and re-suspended in 50 mM Tris pH 7.5, 150 mM NaCl, 1mM EDTA, 1% NP-40, 10% glycerol and incubated on ice for 10 min to lyse mitochondria and other organelles. After spinning for 10 min at 21,000 x g (4°C), the supernatant was stored as mitochondrial extract. Fractions were probed by immunoblot for GAPDH (cytosol), Tomm20 (mitochondria) to determine purity of cellular fractions. Prior to running qPCR, DNA was first extracted from whole cell, cytosolic, or mitochondrial extracts. Each extract was treated with 50 μg/ml RNase A at 37°C for 2 h, then with 200 μg/ml proteinase K at 55°C for 1 hour followed by phenol/chloroform extraction and precipitation. qPCR was performed using PowerTrack SYBR Green master mix with primers for human *ND1*, *ND2* and *D-Loop* (See Appendix). After normalization to nuclear *KCNJ10*, mtDNA abundance in the cytosol was determined relative to whole cell extracts using the delta delta Cq method.

### Immunoblots

Cell monolayers were detached by scraping into ice cold PBS containing 1X Halt^TM^ protease inhibitor cocktail. Pellets were lysed in RIPA buffer (Cell signaling) for 20 min at 4°C and insoluble material pelleted by centrifugation at 16,000 x *g* for 10 min. Supernatants were subject to total protein quantification using a BCA protein assay kit (ThermoFisher) to normalize for protein amount. Lysates were denatured using Bolt^TM^ LDS sample buffer and separated on Bolt^TM^ 8%, 12%, or 4-12% Bis-Tris Plus mini protein gels (according to protein size) using the iBlot2 western blot system (Thermo scientific). PVDF membranes were probed overnight at 4°C with antibodies diluted in 5% BSA in TBST. HRP-conjugated secondary antibodies diluted 1/5000 in 5% skin milk were incubated at RT for 1 hr, and after extensive washing, blots were developed with SuperSignal^TM^ substrate (ThermoFisher) and imaged using an iBright CL1500 imaging system.

### Expression Plasmids and Microscopy

HeLa or THP-1 cells were seeded onto collagen-coated #1.5 round coverslips (ThomasScientific). For THP1 cells, differentiation was achieved by activating with 50 ng/ml PMA (MilliporeSigma) for 24 hr, then rested for 48 hr prior to additional treatment. HeLa cells were transfected with 500 ng of the following plasmids using TransIT®-LT1 Transfection Reagent (Mirus): pEGFP-hGal3 (Addgene #73080) a gift from Tamotsu Yoshimori, pcDNA3-TFAM-mScarlet (Addgene #129573) a gift from Stephen Tait^59^, pGFP-Cytochrome C (Addgene #41181), a gift from Douglas Green^58^, pLamp1-mCherry (Addgene # 45147) a gift from Amy Palmer^93^, pEGFP-C1-APOL3 constructed herein was used for APOL3 imaging experiments in certain cell lines. APOL3 isoform 2 (CCDS13923.1) was used for all experiments since this is the most commonly expressed transcript^76^. To transduce APOL3 into HeLa cells for rescue and imaging experiments, pMSCV-APOL3-eGFP (constructed herein) was used after packaging into retrovirus. To deliver APOL3 into THP-1 cells, APOL3 was cloned into pLVX-APOL3-EGFP (XhoI and XbaI sites) and packaged into lentivirus before selecting with 4 μg/ml puromycin. To image mitochondria via OMP25, Snap-OMP25 (addgene #69599) a gift from David Sabatini ^94^ was packaged into VSV-G pseudotyped lentivirus using HEK293T cells with the packaging plasmid psPAX2. Following transduction into HeLa cells or THP-1 monocytes and plating onto coverslips, cells were incubated with 20 nM JF_646_ SNAP ligand (a gift from Luke Lavis) for 30 min in complete medium. Cells were rinsed in medium then incubated for an additional 10 min to dissipate unbound dye before imaging. When required, transfected or transduced cells were primed with 500 U/ml IFN-γ and 0.1 ng/ml IL-1β (for THP-1) prior to inducing organellar damage. Cells were fixed in 4% w/v paraformaldehyde in PBS (ThermoFisher) for 15 min at RT then rinsed three times with PBS. When imaging transfected cells directly, cells were stained with Hoechst 33258 (ThermoFisher) using a 1/2000 dilution in PBS for 10 min, then mounted onto slides using VectaShield® Plus antifade mounting medium (Vector Laboratories). Slides were allowed to cure for 24 hr prior to imaging.

### Immunofluorescence Microscopy

PFA-fixed coverslips were quenched with 50 mM NH_4_Cl in PBS for 15 min, then permeabilized in 0.2% Triton-X100 (ThermoFisher) for 5 minutes, or 0.1% Saponin (ThermoFisher) for 5 min. For saponin permeabilization, 0.1% saponin was included in all subsequent washes and incubations and removed prior to DAPI staining. Saponin permeabilization was used for all immunostaining involving Lamp1. Permeabilized cells were rinsed in PBST and blocked in 5% BSA in PBST for 2 hr at RT. Coverslips were immunolabeled in blocking buffer overnight at 4°C using the following antibodies: rabbit anti-Tomm20 (Abcam, ab186735, 1/200 dilution), mouse anti-Lamp1 (Invitrogen, MA5-46199, 1/200 dilution), human anti-cardiolipin (USbiological, C1375, 1/500 dilution). After washing 5X with PBST, secondary antibodies conjugated to Alexa fluor^TM^ Plus 488, 594, or 647 (ThermoFisher) diluted to 1 μg/ml in blocking solution were added for 1 hr at RT. After rinsing 5X in PBST, coverslips were stained with Hoechst 33258 (Thermo scientific) using a 1/2000 dilution in PBS for 10 min, then mounted onto slides using VectaShield® Plus antifade mounting medium (Vector Laboratories). Slides were allowed to cure for 24 hr prior to imaging.

### Airyscan super resolution imaging

APOL3^HA^ knock-in cells expressing Snap-OMP25 seeded on collagen-coated #1.5 round coverslips, primed for 18 h with IFN-γ, then stimulated for 2 h with 600 μM LLOME. 20 nM JF_646_ SNAP ligand was added for the final 30 minutes, cells were washed with PBS and fixed with 4% PFA for 20 min at RT. Coverslips were quenched with 50 mM NH_4_Cl in PBS for 15 min, then permeabilized and blocked with 0.1% Saponin in 5% BSA/PBST for 2 hr at RT. Coverslips were immunolabeled in blocking buffer with human anti-cardiolipin (1/500 dilution) and mouse anti-HA.11 epitope tag (BioLegend) (1/200) overnight at 4°C. After washing 5X with PBST, secondary antibodies conjugated to Alexa fluor^TM^ Plus 488, and 594 (ThermoFisher) diluted to 1 μg/ml in blocking solution were added for 1 hr at RT. Slides were mounted using VectaShield® Plus antifade mounting medium (Vector Laboratories) and cured for 24 hr prior to imaging. Slides were imaged using a Zeiss LSM900 with Airyscan 2.

### Mitophagy assay

NTC or APOL3^-/-^ HeLa cells were transfected with pCLBW cox8-EGFP-mCherry (Addgene #78520) a gift from David Chan^50^. 24 hr after transfection, cells were primed with IFN-γ (500 U/ml) for 18 hr, then treated with LLOME (600 μM) or CCCP (10 μM) for 6 hr in the presence of Q-VD-Oph (30 μM) to inhibit cell death and enable visualization of mitophagy events (mCherry positive/GFP negative structures).

### MOMP assays

HeLa cells were transiently transfected with pEGFP-C1-FKBP (Addgene #129575 a gift from Dr. Stephen Tait) and pCDNA3-AIF-mCherry-FRB (Addgene #67530, a gift from Dr. Stephen Tait). After 48 h, cells were for treated with LLOME (600 μM, 2 hr), or ABT-737 (10 μM, 3 hr), in the presence of A/C heterodimerizer (50 nM, Clontech). After fixation with 4% PFA (15 min room temperature), cells displaying MOMP (GFP/mCherry co-localization) were scored using the Leica Thunder Widefield Microscope.

### Magic Red assay

IFN-γ-primed NTC or APOL3^-/-^ cells were incubated for 15 min with Magic Red substrate MR-RR2 (Enzo Life Sciences) to label functional endolysosomes then imaged at specific intervals following a 30-minute pulse with LLOME.

### Image acquisition and analysis

Images were taken using a Leica® Thunder Imager 3D Assay then transferred to ImageJ for analysis. Deconvolution and computational clearing were performed using the Leica® LasX software with the small volume computational clearing algorithm set at 80% strength and optimized for objects 2 μm in diameter. For fractional overlap analysis quantification, images were thresholded to produce regions of interest for compartments positive for each marker. The total pixel area of the intersecting regions was then calculated for each image. For MOMP analysis, mitochondrial volume (MitoCherry) was determined and percentage of pixel area intersecting regions of cytoGFP measured. To count puncta, signal was thresholded to isolate the majority of high intensity puncta. For colocalization analysis, Pearson’s R value was determined for each cell using the Coloc 2 plug-in for ImajeJ.

### Luciferase assays

1 x 10^5^ HeLa cells were seeded into 96 well plates overnight. 9 ng of p561-luc^95^, which harbors IRF3-dependent firefly luciferase consisting of the -134 to +1 of the *IFIT1* gene (provided by S. Ghosh), and 1 ng of pRL-TK (Promega) which drives constitutive expression of *Renilla* luciferase, was transfected into each well using *Trans*IT®-LT1 Trasnfection Reagent (Mirus). 6 h after transfection, cells were primed for 18 h with 500 U/ml IFN-γ. Media was removed for 4 h, cells were pulsed with LLOME (600 μM, 2 hr), then luciferase activity measured using the Dual-Glo® luciferase assay system (Promega). Individual values represent the mean of technical triplicates. Each well was normalized to *Renilla* luciferase to account for differences in cell number and transfection efficiency, and the means of each triplicate then compared with the mean of untreated control triplicate wells that were processed in parallel.

### Immunoprecipitation and qPCR

Human *CGAS* (CCDS4978.1) or *eGFP* was cloned into pMSCV-3XFlag-Puro and used to transduce wildtype or APOL3^-/-^ Hela cells. Immunoprecipitations were performed as previously described^96,97^ with a few modifications. Following selection, approximately 10 x 10^6^ cells were stimulated with 600 μM LLOME for 2 h then rested for 2 h in complete medium before harvesting. When required, Q-VD-Oph (30 μM) was included in the incubation. Cells were lysed in Buffer A containing 20 μg/ml digitonin for 15 min at 4°C and the cytosol clarified by spinning at 21,000 x *g* for 10 min to pellet organelles, then pre-cleared with mouse IgG agarose for 15 min (MilliporeSigma). A 1/10 fraction was removed and defined as input, and the remainder incubated with anti-FLAG® M2 affinity gel (MilliporeSigma) for 2 h with end over end rotation at 4°C. Beads were washed 5X with lysis buffer (Buffer A + Digitonin) for 5 min each then subject to DNA extraction using PicoPure DNA Extraction kit (ThermoFisher). Alternatively, pull-down efficiency was confirmed via immunoblot following elution with 1X Laemmli sample buffer. DNA extracts were analyzed by qPCR for primers specific to mitochondrial (*ND1*, *ND2*) or nuclear (*Line1*, *PolG*, *18S*) transcripts using PowerUP SYBR Master Mix and analyzed using a Bio-Rad CFX96 Real-time PCR system.

### mtDNA depletion

HeLa, THP-1 or primary AECs were cultured on 96 well plates coated with rat tail collagen (MilliporeSigma). Initial cell concentrations for HeLa cells were 5 x 10^3^ cells/well, for THP-1 7.5 x 10^3^ cells/well and for primary AECs 1 x 10^4^ cells/well. Cells were grown for 72 to 96 hr (HeLa or THP1) or for 144 hr (AECs) in complete medium containing 80 μg/ml 2′,3′-dideoxycytidine (ddC) (MilliporeSigma). Control cells were treated the same way but without ddC. Once the cells reached 75-85% confluence as determined using the Millcell® Digital Cell Imager (MilliporeSigma), experiments were performed. Duplicate wells were subject to RNA extraction and qPCR to determine mtDNA depletion efficiency using primers for *ND1* and normalizing to nuclear DNA using primers against *KCNJ10*.

### rAPOL3 protein purification

APOL3 was purified from *E. coli* inclusion bodies as previously described^36^. Briefly, Overnight cultures of BL21(DE3) pLysS harboring the APOL3 expression plasmid (pET28a-6XHis-PP) were grown in LB containing kanamycin (50 μg/ml) and chloramphenicol (20 μg/ml). Cultures were grown to OD_600_ = 0.7 in media without chloramphenicol and induced with 1 mM IPTG for 4 hours at 37°C. Cell pellets were lysed in 50 mM Tris pH 8.0, 5 mM EDTA and lysozyme (100 μg/ml; Sigma) with sonication. Insoluble material was pelleted at 20,000 x *g* and washed with lysis buffer containing 0.5 M NaCl. Pellets were solubilized in 6 M guanidine hydrochloride, 50 mM potassium phosphate pH 8.0, 1 mM TCEP, 10 mM imidazole for 1 hour at room temperature and protein was affinity purified using Ni-NTA beads (Qiagen) and dialyzed extensively into 20 mM acetic acid. For most experiments, the His-tag fusion protein was digested with His-3C protease (Genscript) in 50 mM MES pH 6.5 overnight at 4°C. To remove the tag, undigested protein and the protease, the reaction and all insoluble precipitate was solubilized in 6 M guanidine hydrochloride, 50 mM potassium phosphate pH 8.0, 1 mM TCEP, 10 mM imidazole and incubated overnight with fresh Ni-NTA beads. Flow through was collected and dialyzed extensively into 10 mM acetic acid and flash frozen. For experiments in which rAPOL3 was pulled down from mitochondrial reactions, His-rAPOL3 was used without incubating with 3C protease to remove the tag.

### Liposome preparation and leakage experiments

Liposomes were prepared as described previously^36^. Phospholipids (POPC, POPE, PI, CL) at a ratio of 60:30:10 in chloroform were mixed and evaporated overnight under vacuum. Lipid film was hydrated in 50 mM MES pH 6.5, 100 mM potassium gluconate (KGl) and 80 mM calcein. Lipids were solubilized with continual vortexing followed by five freeze/thaw cycles and then extruded through a 0.1 μm polycarbonate filter (Avanti Polar Lipids Inc.) 40 times using a mini-extruder device (Avanti Polar Lipid Inc.). Non-encapsulated calcein was removed using Illustra Microspin G50 columns (GE Health care). To measure leakage, liposomes of the indicated composition (500 μM lipid) were mixed with rAPOL3 (1 μM) in lipid solubilization buffer and fluorescence measured using a BioTek multi-mode Cytation plate reader with excitation and emission wavelengths λ_Ex_ = 495 nm and λ_Em_ = 525 nm for calcein. Fluorescence prior to addition of protein was treated as *F*_t0_. 5 μl of rAPOL3 diluted in 10 mM acetic acid was added after ∼1 min, and fluorescence recorded continuously (at 2 s intervals). 5 μl of final dialysate (20 mM acetic acid) was used as a mock treatment. 10 μl of 1% Triton X-100 was added to achieve complete dye release and the average of the maximal value defined as *F*_t100_. The percentage of dye efflux at each time point was calculated as (*t*) (%) = (*F*_t_ – *F*_t0_) × 100/(*F*_t100_ – *F*_t0_).

### Fluorescent lipid experiments

100 nm POPC/POPE/CL Liposomes were prepared as described above with the following modification: 10% TopFluor^TM^-labeled cardiolipin (Avanti) was used as the source of CL (10% of total lipid), or TopFluor^TM^ PE was spiked (30% of total PE) into the POPE to achieve an equivalent final concentration as CL (10% total lipid). Unincorporated lipids were removed by Microspin G50 columns (GE Health care). Fluorescence intensity (λ_Ex_ = 498 nm and λ_Em_ = 507) nm of the liposomes in solution was measured using a Biotek Cytation multi-mode plate reader and defined as *F*_t0_. 6His-rAPOL3 was added to a final concentration of 2 μM in incubation buffer (50 mM MES pH 6.5, 100 mM KGl, 50 mM NaCl) for thirty minutes (with frequent agitation) before separating the soluble and insoluble components by centrifugation at 10,000 x *g* for 15 min (RT). rAPOL3 lipoprotein complexes were recovered from the soluble fraction by first binding to Ni-NTA agarose in incubation buffer containing 25 mM imadozole to reduce non-specific binding. Agarose was washed 5X in the same buffer then eluted with 400 mM imidazole. Fractions containing protein (estimated by A280) were pooled, concentrated to roughly the same volume as input, and the fluorescence measured as *F*_t_. Percentage of lipid extracted/solubilized by rAPOL3 was calculated as *F*_t_/*F*_t0_ x 100.

### Isolation and treatment of mitochondria

HeLa cells in confluent T 175 flasks were collected by gentle scraping and pellets resuspended in Buffer A and homogenized with a glass Dounce homogenizer. 1 mM EGTA and 2 μM rotenone (MilliporeSigma) were included in all buffers to safeguard mitochondrial integrity. Nuclear fraction was removed by centrifugation at 1000 x *g* for 10 min at 4°C. Supernatant was decanted and recentrifuged at 6000 x *g* for 10 min at 4°C. Mitochondrial pellets were gently resuspended in Buffer A lacking EGTA/rotenone lacking EGTA/rotenone and used within 2 h. Freshly isolated mitochondria (1 mg/ml) were incubated with recombinant GST-BAX (10 μg/ml) (MilliporeSigma), rAPOL3 (10 μg/ml diluted in 10 mM acetic acid), or Fe(II)SO_4_ (50 μM)/ascorbate (500 μM) for 30 min at 37°C with constant shaking. 1% NP-40 (EMD Millipore) was used to release total mtDNA. Treated mitochondria were pelleted (10,000 x *g*, 15 min, 4°C) and the supernatant collected. dsDNA was quantified in the supernatants using Quant-iT^TM^ PicoGreen^TM^ dsDNA assay kit and normalized to total mtDNA released by NP-40 treatment. Cardiolipin content in the recovered supernatant was measured using the cardiolipin assay kit (Abcam, ab241036) according to manufacturer’s recommendation. In some experiments, supernatants were processed further to isolate rAPOL3 lipid protein complexes from completed mitochondrial reactions. For this, supernatants were added to Ni-NTA agarose and subject to end-over-end incubation for 1 hr at RT. Agarose was washed 5X with Buffer A containing 200 mM NaCl in the place of sucrose and 20 mM imidazole. rAPOL3 was eluted across an imidazole gradient (50 to 500 mM) and fractions were collected, concentrated, and analyzed for cardiolipin content (cardiolipin assay kit, Abcam) and by SDS-PAGE followed by Coomassie Brilliant Blue-R250 staining. Size exclusion chromatography (Superdex 200 Increase; GE Healthcare) equilibrated with 10 mM Tris-HCl pH 7.0, 100 mM KGl, 50 mM KCl containing 10 mM imidazole was used to analyze pooled eluates from Ni-NTA purifications.

### ELISAs

IFN-β was quantified clarified cell supernatants using the Human IFN-beta Quantikine ELISA (R&D systems) or Human Interferon beta ELISA kit (Abcam) and IL-1β detected by the Human IL-1 beta/IL-1F2 ELISA Kit (R&D systems) as described by the manufacturer.

### Cell lines and treatment conditions

HeLa (CCL2), 293T/17 (CRL-1573) cells were purchased from ATCC. Cells were grown in DMEM supplemented with 10% (v/v) heat-inactivated fetal bovine serum (FBS) at 37°C in a 5% CO_2_ incubator without antibiotics. THP-1 (TIB-202) cells were a gift from Donna Farber and were grown in RPMI supplemented with 10% FBS. Cells were routinely tested for mycoplasma contamination using Lookout® One-Step Mycoplasma PCR detection kit (Millipore Sigma). Where indicated, cells were pre-treated with 500 U/ml recombinant IFN-γ (PBL Assay Science) for 18 hr, with both 500 U/ml IFN-γ and 0.1 ng/ml recombinant IL-1β (ThermoFisher) then treated with the designated stimulus in media lacking cytokines Concentrations are as follows: LLOME (Millipore Sigma) was used at 600 μM unless otherwise indicated, Silica nanoparticles (Invivogen) suspended in ultrapure water and used at 50 μg/ml, Streptolysin O (Millipore Sigma) used at 2 U/ml, Cholera toxin B subunit (Millipore Sigma) was used at 10 μg/ml, CCCP (Millipore Sigma) was used at 10 μM, ABT-737 (MedChemExpress) was used at 2 μM, 5 μM, or 10 μM, Q-VD-OPh (MedChemExpress) was used at 30 μM, Poly(I:C) (Invivogen) was used at 2 μg/ml, Mdivi-1 (HY-15886; MedChemExpress) was used at 10 μM, urolithin A (HY-100599; MedChemExpress) was used at 2 μM, TiO_2_ particles (<25 nm in diameter; MilliporeSigma) was used at 50 μg/ml.

### Primary cell isolation

Whole lung tissue was obtained from adult brain-dead organ donors through an approved protocol and material transfer agreement with LIVEOnNY, the local organ procurement organization for the New York Metropolitan area as described previously^98^. Tissues selected for this study were listed as non-smokers, had no history of asthma and were negative for SARS-CoV-2, cancer, hepatitis B, and C, and human immunodeficiency virus. This study is not human subject research because the tissues were obtained from deceased individuals, as confirmed by the Institutional review board at Columbia University. Lung tissue was subject to mechanical and enzymatic digestion, followed by density centrifugation as previously described^99^. Lymphocytes were removed by EasySep^TM^ CD45 depletion kit II (STEMCELL Technologies)) and airway epithelial cells (AECs) isolated using EasySep™ Human EpCAM Positive Selection Kit II (STEMCELL Technologies) according to manufacturer’s instructions. AECs were cultured in BEGM™ BulletKit™(Lonza) and used within 6 passages. Following lentiviral transduction with Cas9/sgRNA expressing lentivirus, selection was achieved with 2 μg/ml puromycin for 72h before overnight treatment with IFN-γ (500 U/ml) and addition of 50 μg/ml TiO_2_ particles (<25 nm in diameter) (Sigma) for 24 hr.

### Detection of mitochondrial membrane potential

Primary human AECs from three different donors were primed (or not) with 500 U/ml IFN-γ for 18 hr then treated with 50 μg/ml TiO_2_ for 12 hr. Cells were washed with warm Hank’s Balanced Salt Solution (HBSS) and incubated with 20 nM TMRM dye (ThermoFisher) in complete medium for 30 min at 37°C, 5% CO_2_. Cells were again washed with warm HBSS, detached with TrypLE reagent (ThermoFisher) and analyzed by flow cytometry using a LSR Fortessa, LSR II Flow cytometer. Median Fluorescence Intensity (MFI) was determined from three donors, each done in technical triplicate.

### Infections

*Salmonella enterica* serovar Typhimurium (*Stm*) strain 1344 (a gift from J. Galan), Δ*sifA* mutant (a gift from J. Brumell)^100^ were used for infections. For *Salmonella* infections in SPI-1 conditions, overnight bacterial cultures were diluted 1:33 in LB, grown for 3 hours (37°C shaking at 300 rpm) washed twice in PBS and used to infect cells at 80% confluence at an MOI of 10 unless otherwise indicated. Plates were centrifuged for 10 min at 1000 x g (room temperature) and incubated for 1 h at 37°C to allow invasion. Extracellular bacteria were killed by replacing inoculum with fresh DMEM containing 100 μg/ml Gentamicin. For SPI-2 conditions, cultures were incubated overnight at 37°C without shaking, washed with PBS and used to infect cells directly. Recombinant adenovirus was prepared using the pAdEasy system as described^101^ and enlisted the pAdTrack shuttle vector which is E1/E3 deleted (Addgene #16404, a gift from Bert Vogelstein^102^). Virus was prorogated in HEK293 (ATCC CRL-1573) and purified using double CsCl_2_-banding. The P137L mutation was introduced using the Phusion site-directed mutagenesis kit (ThermoScientific). Copy number was estimated according to the OD_260_ method where 1 OD_260_ = 1.16 x 10^12^particles/ml^103^. Infections were performed in 12 well plates with 100 virus particles/cell in 100 μl for 2 hours at 37°C with gentle rocking. Inoculum was removed and replaced with fresh media for the duration. AdV-*ts1* were prorogated at permissive temperature (33.5°C), before conducting experiments at non-permissive temperature (38.5°C)^104^.

### Statistics

No sample size calculation or blinding was performed. For quantification of micrographs, sample size reflects prior knowledge of variation and the number of events that could reasonably be quantified. Samples were allocated into experimental groups randomly. Data were analyzed by Graphpad Prism 10.0 software. Unless indicated otherwise, statistical significance was determined by *t* test (two-tailed) or one-way analysis of variance (ANOVA) with Tukey’s multiple-comparison test.

