## Supplemental information for "The antibacterial factor APOL3 couples lysosomal damage to mitochondrial DNA efflux and type I IFN induction"

### Figure S1

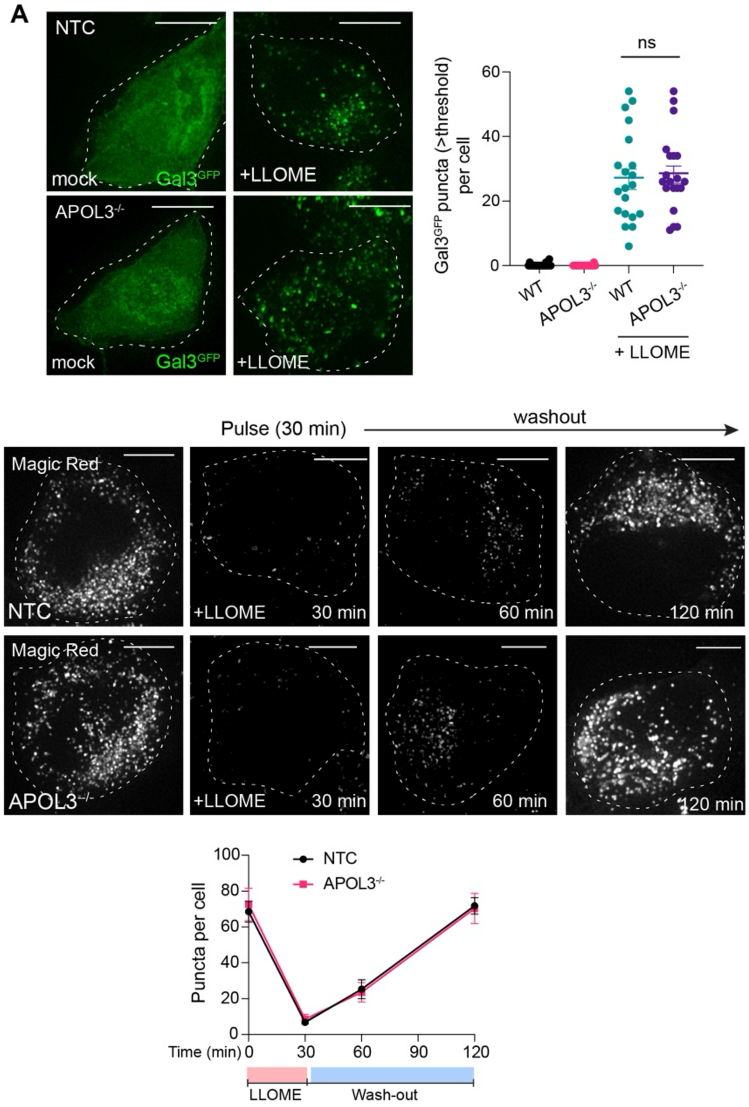


**Figure S1. Deletion of APOL3 does not affect acute sensitivity to, or recovery from, transient lysosomal damage.**

**(A)** IFN-γ-primed HeLa genotypes expressing Galectin-3-GFP (Gal3$^{GFP}$) were treated with LLOME (30 min) and analyzed by microscopy. Amount of GFP puncta per cell ($n$ = 20 cells, mean ± SD) is quantified. Shown are single *z* planes of Thunder de-convolved images (scale bar, 10 μm). ns (non-significance following student *t* test). **(B)** IFN-γ-primed HeLa cells of the indicated genotype were incubated with Magic Red to label functional lysosomes then imaged following a 30-minute pulse of LLOME. Shown are maximum intensity projections (scale bar, 5 μm). Magic Red puncta per cell was quantified from 15 cells across 4 biological replicates (mean ± SEM).

### Figure S2

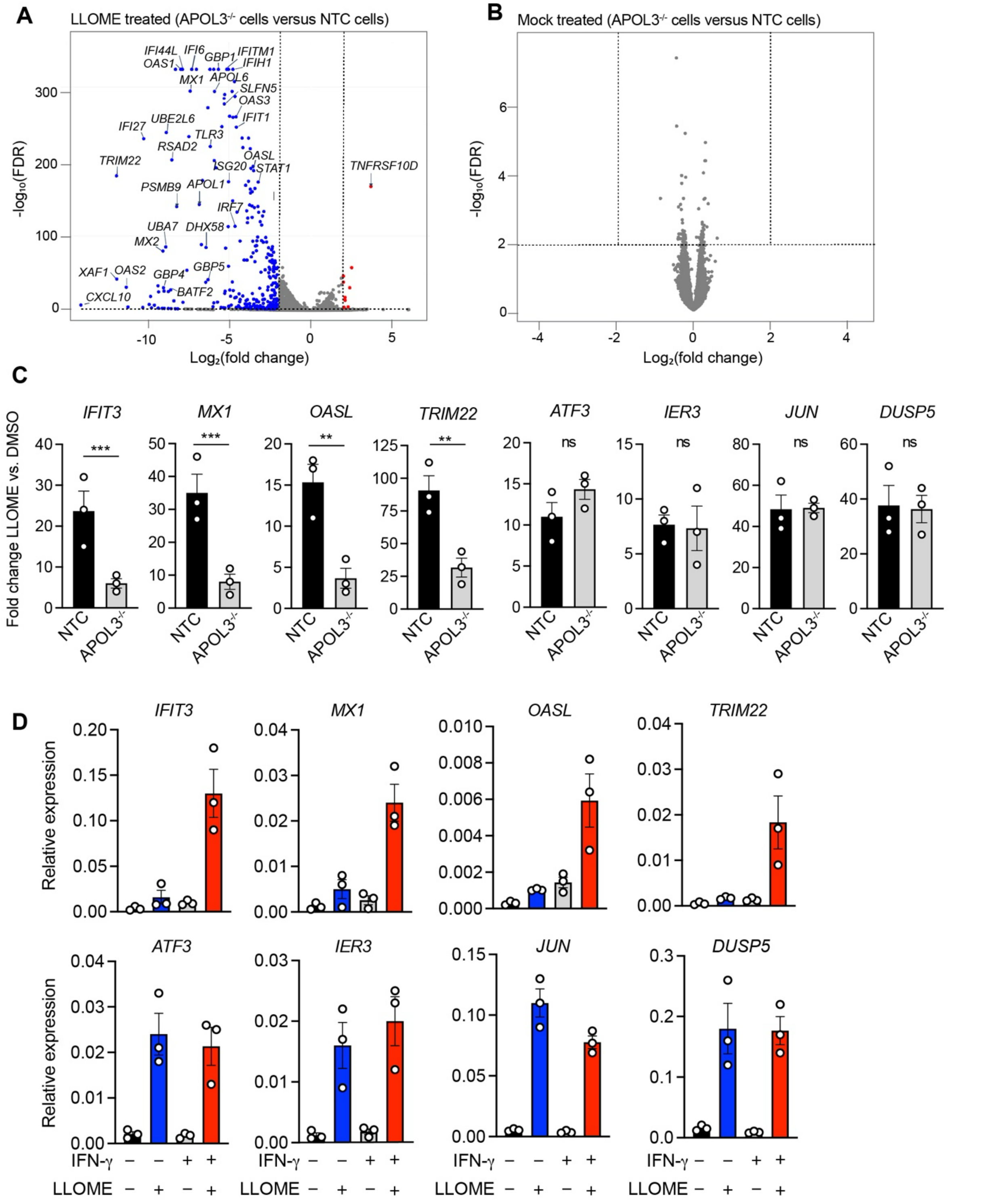


**Figure S2. IFN-γ-stimulated APOL3 is required for upregulation of ISGs but not stress response genes during lysosomal damage.**
**(A, B)** Volcano plots showing differential expression of LLOME-treated (A) or untreated (B) NTC cells compared to APOL3$^{-/-}$ cells. Significantly upregulated genes ($P < 0.01$, log-fold change (LFC) $> 2$) are

depicted in red, and significantly downregulated genes ($P < 0.01$, log-fold change (LFC) > 2) are depicted in blue. Selected genes of interest were annotated. Statistical significance for 2 independent biological replicates was Statistical significance was calculated using the Wald test, with P values adjusted by the Benjamini-Hochberg procedure. **(C)** IFN-γ-primed NTC or $APOL3^{-/-}$ cells were stimulated with LLOME or vehicle (DMSO) for 2 h, allowed to recover for 8 h, then analyzed by qPCR for the indicated transcript. Data is relative fold change in expression after LLOME treatment compared with DMSO control and represents the mean ± SEM from 3 independent experiments. ** $P < 0.01$; *** $P < 0.001$ determined by two-tailed Student's t test. ns, not significant. **(D)** qPCR assessment of wildtype (parental) HeLa cells with or without IFN-γ-priming treated with LLOME (or DMSO) as in (C). Gene expression is normalized to *GAPDH* and represents mean ± SEM from 3 independent experiments.

### Figure S3

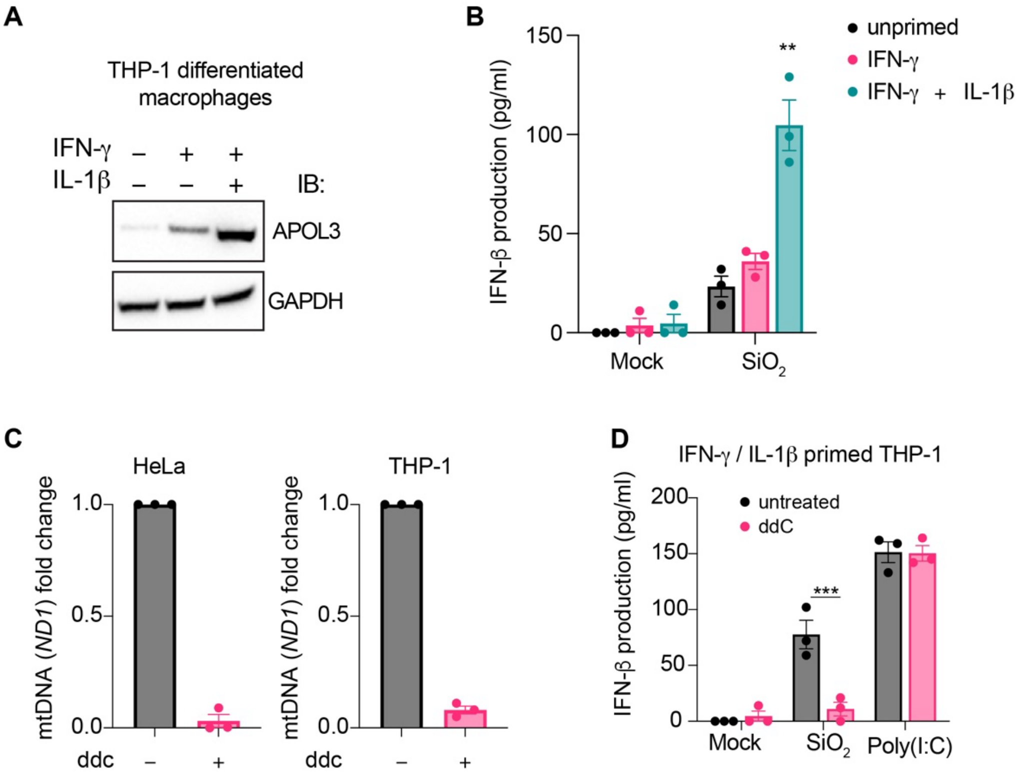


**Figure S3**. **Phagosomal disruption elicits mtDNA-dependent type I IFN from IFN-γ / IL-1β-primed THP-1 macrophages.**

**(A)** Immunoblot (IB) for APOL3 or GAPDH in cell lysates prepared from THP-1 differentiated macrophages with overnight priming with the cytokines IFN-γ and/or IL-1β. **(B)** IFN-β protein levels surveyed by ELISA in the supernatants of primed or unprimed THP-1 macrophages 24 hr after being fed $SiO_2$ particles (50 μg/ml for 4 hr). ** P < 0.01, determined by one-way ANOVA Tukey's multiple comparison test. **(C)** qPCR assessment of whole-cell mtDNA (*ND1* transcript) in HeLa cells or THP-1 macrophages cultured for 96 hr with or mtDNA depletion using 2',3'-dideoxycytidine (ddC) (80 µg/ml). **(D)** IFN-β protein levels in the supernatants of IFN-γ / IL-1β. primed THP-1 macrophages 24 hr after being fed $SiO_2$ particles (50 μg/ml for 4 hr) or treated with Poly(I:C) (2 µg/ml for 24 hr). Cells were cultured with or without ddC for 96 hr to deplete mtDNA prior to treatment. *** P < 0.001 determined by two-tailed Student's t test. Immunoblots (A) are representative of 2 independent experiments and results in (B), (C), (D) are mean ± SEM from 3 independent experiments

### Figure S4

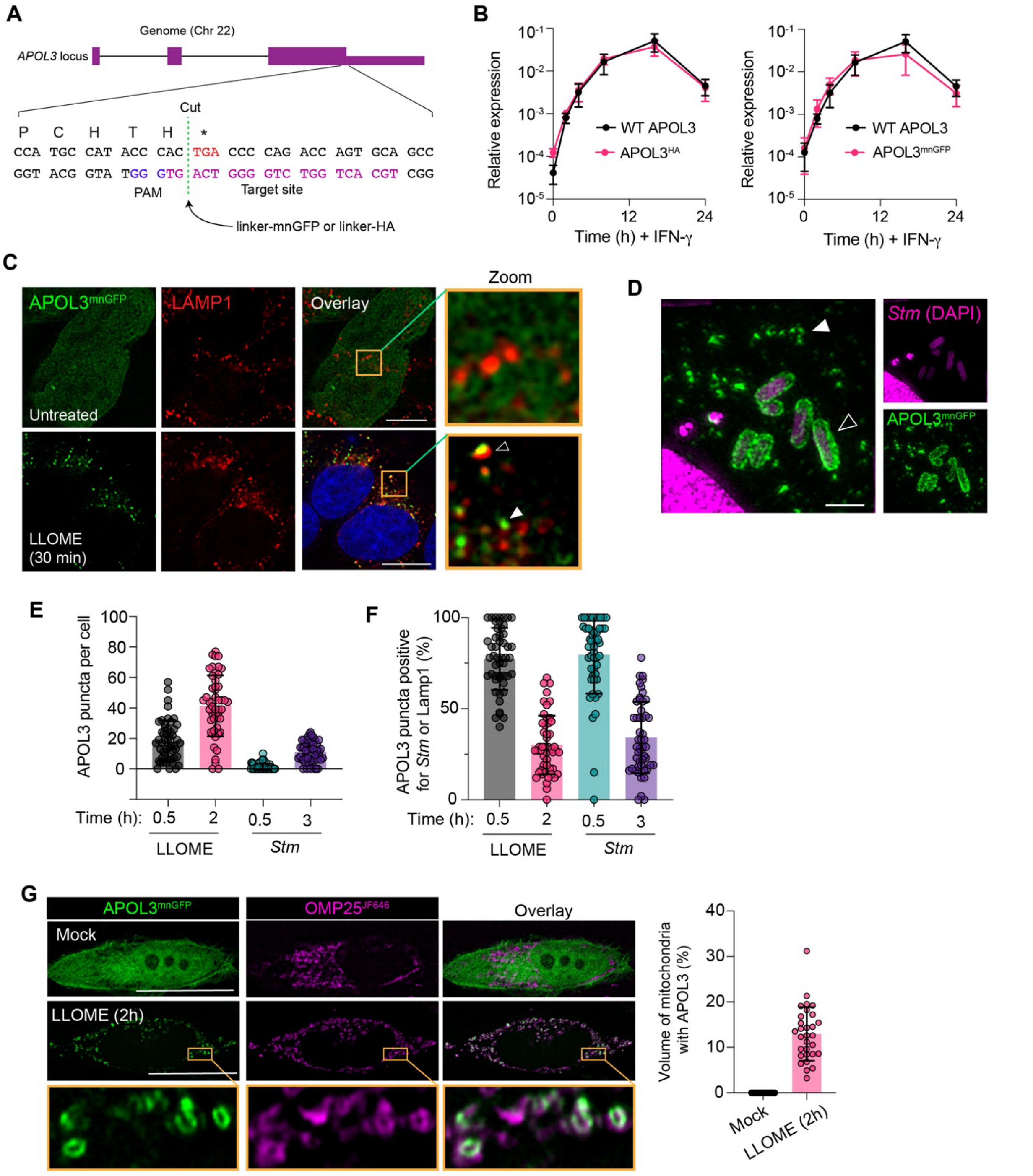


**Figure S4. Endogenous APOL3 targets mitochondria during sterile and infectious lysosomal damage.**
**(A)** Schematic depicting construction of APOL3 knock-in cells using the CRISPaint gene tagging system. **(B)** qPCR analysis of *APOL3* induction upon IFN-γ treatment before or after gene tagging. Parental *APOL3* (purple) is upregulated to a similar degree as tagged APOL3$^{mnGFP}$ and APOL3$^{HA}$. Results are presented as expression relative to *GAPDH* and are mean ± SEM from 3 independent experiments. **(C)** IFN-γ-primed

$APOL3^{mnGFP}$ HeLa cells were treated with LLOME (30 min) and immunostained for Lamp1. Shown are single *z* slices from Thunder deconvolved widefield images. Filled white arrow depicts Lamp1-negative APOL3 foci, outlined arrow depicts Lamp1-positive APOL3 foci. **(D)** IFN-γ-primed $APOL3^{mnGFP}$ HeLa cells infected with *Salmonellla enterica* serovar Typhimurium (*Stm*) for 2 h and subject to Airyscan super-resolution imaging. *Stm* is marked by DNA (DAPI) staining. Filled white arrow depicts *Stm*-negative APOL3 foci; outlined arrow depicts *Stm* positive for APOL3. Shown are maximum intensity projections. **(E, F)** IFN-γ-primed $APOL3^{mnGFP}$ HeLa cells were treated with LLOME as in (C) or infected with *Stm* as in (D). The total number of APOL3 puncta per cell at the indicated time was quantified in (E), and the percentage of APOL3 puncta that overlap with Lamp1 (lysosomes) or DAPI (*Stm*) is quantified in (F). Shown is mean $\pm$ SD from 50 cells analyzed per condition. **(G)** IFN-γ-primed $APOL3^{mnGFP}$ HeLa cells expressing lentiviral-delivered SNAP-tag OMP25 ($OMP25^{JF646}$) were treated with LLOME for 2 hr and imaged. Volume of total OMP25 signal that overlaps with APOL3 foci is quantified for 30 cells (mean $\pm$ SD). Images and results in (E) (F) and (G) are representative of 3 independent experiments. Scale bars, 5 μm (C), 2 μm (D), 10 μm (G).

### Figure S5

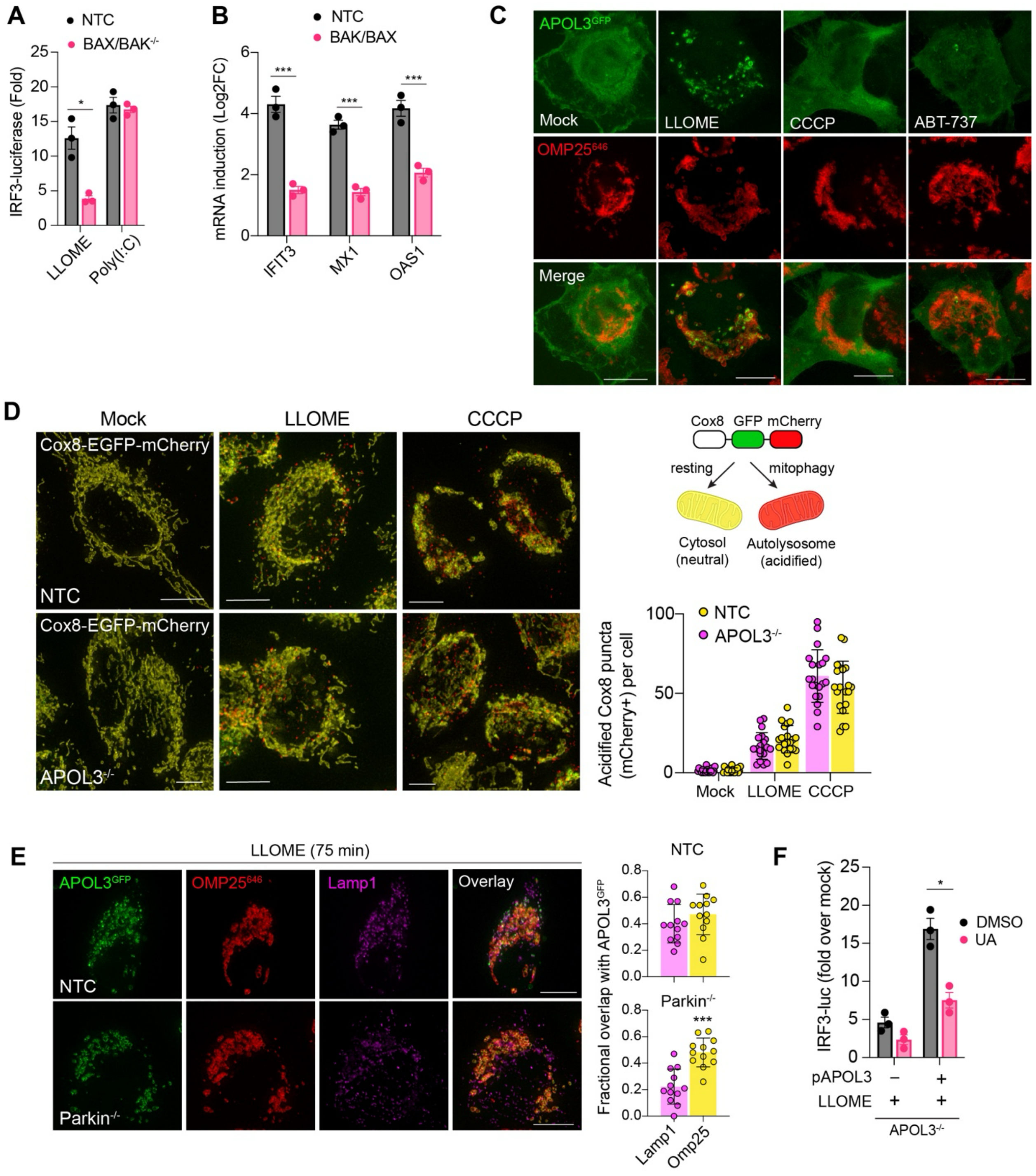


**Figure S5. Mitochondrial damage is required but not sufficient to recruit APOL3 which is independent of mitophagy.**

**(A)** IFN-γ-primed NTC or BAX/BAK$^{-/-}$ HeLa cells expressing IRF3-luciferase were pulsed with LLOME (600 μM, 2 hr) or treated with Poly(I:C) (2 μg/ml) and luminescence determined after 6 hr. Results are fold change relative to mock treated. **(B)** IFN-γ-primed NTC or BAX/BAK$^{-/-}$ HeLa cells were pulsed with LLOME for 2 hr and analyzed by qPCR for the indicated genes after an additional 4 hr recovery. Results are log2-fold change (Log2FC) relative to mock treated. **(C)** IFN-γ-primed HeLa cells expressing APOL3-GFP and $OMP25^{646}$ were stimulated with LLOME (600 μM, 2 hr), CCCP (10 μM, 2 hr), or ABT-737 (10 μM, 3 hr). Shown are maximum intensity projections from Thunder deconvolved widefield images. **(D)** IFN-γ-primed NTC or $APOL3^{-/-}$ HeLa cells expressing Cox8-EGFP-mCherry were treated with LLOME (600 μM) or CCCP (10 μM) for 6 hr in the presence of Q-VD-Oph (30 μM) to inhibit cell death and enable visualization of mitophagy events. At neutral pH, mitochondria express both eGFP and mCherry and appear yellow. Within acidified compartments only mCherry retains fluorescence and signify mitochondria within autophagosomes. Images are maximum intensity projections of Thunder de-convolved *z* stacks. Quantification depicts number of mCherry aggregates per cell (mean ± SD, n = 20 cells), representative of 3 independent experiments. **(E)** IFN-γ-primed NTC or $Parkin^{-/-}$ HeLa cells expressing APOL3-GFP and $OMP25^{646}$ were treated with LLOME (600 μM) for 75 min. Lysosomes were marked by Lamp1 immunostaining. Representative images are maximum intensity projections of Thunder de-convolved *z* stacks. Fractional overlap is shown for APOL3 with Lamp1 or OMP25 (n = 12 cells, representative of 3 independent experiments, error bars ± SD). **(F)** IFN-γ-primed $APOL3^{-/-}$ HeLa cells complemented with pMSCV-APOL3 or empty pMSCV and transfected with IRF3-luciferase were pulsed with LLOME for 2 hr then luminescence determined after 6 hr in the presence or absence of the mitophagy inducer urothilin A (20 μM). (A), (B) and (F) are mean ± SEM from three independent experiments. *P < 0.05; ** P <0.001 determined by two-tailed Students *t*-test (E), or one-way ANOVA with Tukeky's multiple comparison test (A, B). Scale bar, 5 μm.

### Figure S6

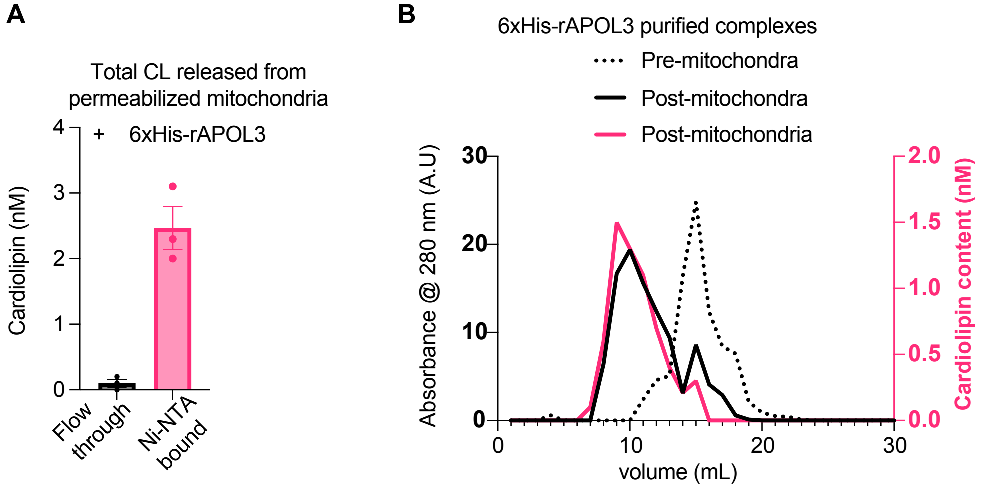


**Figure S6. Cardiolipin (CL) released from permeabilized mitochondria is contained within APOL3-lipoprotein complexes**

**(A)** Mitochondria purified from HeLa cells were incubated with 6xHis-rAPOL3 and rBAX for 30 min. Supernatant was collected after centrifugation and subject to Ni-NTA purification to capture His-tagged proteins. Cardiolipin content of the eluate (bound) or flow-through (unbound) was determined by fluoregenic assay. Data is mean ± SEM from 3 independent experiments. **(B)** 6xHis-rAPOL3 analyzed by size exclusion chromatography before (dashed line) and after incubation with rBAX and mitochondria (solid lines) as in (A). Cardiolipin content (rose) of each elution fraction was determined by fluoregenic assay and protein content (black) estimated by absorbance at 280 nm. Data is representative of 2 independent experiments.

### Figure S7

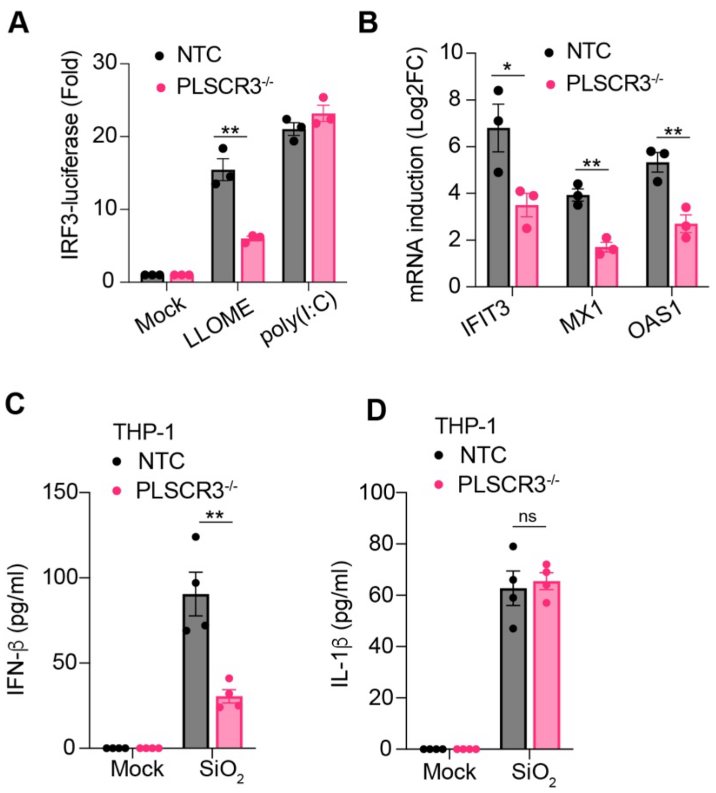


**Figure S7. PLSCR3 is required for the type I IFN response to transient lysosomal damage.** **(A)** HeLa genotypes expressing IRF3-driven luciferase pulsed with LLOME (2 hr) or treated continuously with Poly(I:C) and luminescence determined after 6 hr. **(B)** IFN-γ-primed NTC or PLSCR3$^{-/-}$ HeLa cells were pulsed with LLOME for 2 hr and analyzed by qPCR for the indicated genes after an additional 4 hr recovery. Results are log2-fold change (Log2FC) relative to mock treated. **(C**, **D)** ELISA measurements of IFN-β or IL-1β in the supernatants of IFN-γ / IL-1β-primed THP-1 macrophages of the indicated genotype 24 hr after being fed $SiO_2$ nanoparticles (50 μg/ml for 4 hr) to induce phagosomal rupture. (A) to (D) represent the mean ± SEM from three or four independent experiments. *$P < 0.05$; ** $P < 0.001$ determined by one-way ANOVA with Tukey's multiple comparison test (A) and (B) or two-tailed Students *t*-test (C) and (D).
